# Organ-resolved endothelial regulatory programs link aging and metabolic overload to vascular immune remodeling

**DOI:** 10.64898/2026.07.02.736039

**Authors:** Masataka Yokoyama, Akitoshi Nakayama, Yuki Taki, Mingyang Chen, Yingbo Gong, Moeko Shiina, Takashi Kono, Masanori Fujimoto, Kaoru Ito, Jun-ichiro Ikeda, Tomoaki Tanaka

**Affiliations:** Department of Molecular Diagnosis, Chiba University Graduate School of Medicine, Chiba University, Chiba 260-8670, Japan; Research Institute of Disaster Medicine, Chiba University, Chiba 260-8670, Japan; Beth Israel Deaconess Medical Center Division of Endocrinology, Diabetes & Metabolism, Boston, MA 02115, United States; Department of Advanced Biomedical Data Science, Graduate School of Medicine, Chiba University, Chiba 260-8670, Japan; Department of Diagnostic Pathology, Graduate School of Medicine, Chiba University, Chiba 260-8670, Japan

## Abstract

Systemic aging and metabolic overload remodel the vasculature; however, how endothelial cells integrate these stresses across organs remains unclear. Using multi-organ single-cell and spatial transcriptomics with functional validation, we mapped endothelial and hematopoietic responses in adipose tissue, skeletal muscle, liver, and heart. Organ-specific endothelial transcriptional features were relatively preserved, whereas chronic stress selectively reconfigured regulatory programs: aging induced a conserved *Irf/Stat*-centered endothelial program, while high-fat diet engaged organ-biased lipid and remodeling programs. Spatial analysis revealed perivascular niches centered on aging-associated interferon-stimulated endothelial activation, with neighboring immune and stromal cells expressing C3 and LRP1-associated signals. Rather than simply amplifying inflammation, these niches contained mechanisms that restrained IFN activation, as C3 depletion upregulated vascular IRF7 expression. In parallel, the IFN downstream effector BST2 promoted anti-inflammatory macrophage differentiation and suppressed atherosclerosis. These findings define vascular inflammaging as an organ-resolved niche process in which endothelial IFN activation is coupled to local inflammatory restraint.

**Highlights:**

- Aging induces a shared endothelial type I IFN program across organs.
- A high-fat diet triggers organ-biased endothelial remodeling programs.
- Perivascular interferon niches couple inflammation with local restraint.
- IFN-induced endothelial BST2 promotes CD200R-associated macrophage regulatory features.

## Introduction

Endothelial cells (ECs) form a dynamic interface between the circulation and parenchymal tissues, coordinating vascular development, organ maintenance, and regeneration^1–4^. They act as transcriptionally specialized signaling hubs that release angiocrine factors and adapt vascular function to local tissue demands^5^. Advances in cell type-resolved and single-cell transcriptomic technologies have further established that ECs possess distinct molecular identities across organs and vascular beds^6^. Subsequent studies have defined specialized endothelial programs in individual tissues, including hepatic sinusoidal differentiation, brain vascular zonation associated with blood–brain barrier specialization, kidney vascular specialization, and liver endothelial maturation during development and regeneration^7–10^.

These organ-specific observations were later integrated into a whole-body view by single-cell vascular atlases, which revealed extensive endothelial heterogeneity across tissues and vascular beds^11^. Subsequent human endothelial atlases further defined the transcriptomic signatures of organ- and vascular-bed-specific ECs across multiple tissues^12,13^. More recently, human vascular cell atlases have extended this framework by defining organotypic vascular cell states across multiple human organs^14^. Emerging human aging EC atlas resources have also begun to describe age-associated endothelial transcriptional changes across diverse tissues^15^. Aging-associated vascular remodeling has been further explored using multi-organ single-cell atlases, including Tabula Muris Senis, which demonstrated broad age-dependent transcriptional changes across mouse tissues^16^. Despite this progress, most existing atlases remain primarily descriptive. They have established that endothelial identity differs by organ and changes with age, but have not resolved how ECs across organs respond to chronic systemic stress through shared versus organ-specific transcriptional regulatory programs.

Systemic stresses such as aging, metabolic overload, and chronic inflammation act throughout the body, yet vascular disease develops with marked organ selectivity, from coronary and peripheral atherosclerosis to metabolic remodeling of the hepatic and adipose vasculature. Although hemodynamic forces and vascular geometry contribute to this selectivity, organ-specific endothelial regulatory programs and immune–stromal communication may shape how each tissue interprets the same stress. Single-cell profiling under obesity conditions has revealed organ-specific endothelial vulnerabilities with inflammatory and metabolic remodeling^17^, and spatial analysis of the aging heart has revealed vessel-associated niches as hotspots of cardiac aging and inflammaging^1^. However, whether stress-induced endothelial inflammation represents a uniform systemic response or is reshaped by organ-specific regulatory networks and local immune-stromal niches, and how these programs amplify or restrain vascular inflammation remain unclear.

To address this gap, we profiled endothelial responses across adipose tissue, skeletal muscle, liver, and heart under aging conditions and high-fat diet stress, using single-cell transcriptomics to elucidate organ-specific endothelial activation states and the transcription factor networks that regulate them. This regulon-based framework revealed that aging induces a shared endothelial type I IFN program associated with Irf/Stat regulons across organs, whereas metabolic stress engages more organ-biased lipid metabolic and tissue remodeling programs. By projecting these programs onto tissue architecture, we found that aging-associated interferon-stimulated genes form perivascular inflammatory niches. Further analysis of endothelial–immune communication revealed counterregulatory mechanisms within these niches, including C3-associated modulation of perivascular IFN inflammation and a heart and skeletal muscle-enriched endothelial BST2 axis that restrains macrophage inflammatory propagation and atherosclerotic lesion formation. Our findings suggest that the niche-directed modulation of endothelial–immune communication may enable more precise intervention in vascular aging and disease.

## Results

### Organ-specific gene expression and transcriptional regulation in endothelial cells

To compare endothelial responses to aging and metabolic stress across organs, the mice were assigned to normal diet or high-fat diet (HFD) groups and maintained for either 3 months or 1 year (Figure 1A). Both aging and HFD loading increased body weight and blood glucose levels, confirming the systemic physiological effects of these interventions (Figure S1A). Vascular ECs were isolated from adipose tissue, skeletal muscle, liver, and heart, which represent metabolically active organs with distinct vascular functions. Because endothelial stress responses are closely linked to inflammatory remodeling, hematopoietic cells were collected from the same tissues to enable parallel analysis of ECs and immune cell populations (Figure 1A). After quality control, single-cell RNA sequencing yielded 100,730 cells. UMAP analysis separated the vascular, stromal, and hematopoietic compartments (Figure 1B). Cells were distributed across all four organs, allowing cross-organ comparisons of endothelial and immune compartments (Figure S1B). Vascular cells were identified by *Pecam1* and *Cdh5* expression and were clearly separated from *Ptprc*-positive hematopoietic cells and *Col1a1*/*Pdgfra*-positive fibroblasts (Figures 1B, S1C, and S1D). Feature plots of representative marker genes further confirmed the separation of major vascular, stromal, and hematopoietic populations (Figure S1E). Hematopoietic cells were annotated into macrophage/monocyte/DC populations (Macrophage lineage), T cell/NK cell/ILC populations (T-cell lineage), B cell/plasma cell populations (B-cell lineage), neutrophils, and mast cells using canonical lineage markers, including *Cd68* and *Csf1r* for the Macrophage lineage, *Cd3e*, *Cd3d*, and *Nkg7* for the T-cell lineage, *Ms4a1* and *Cd19* for the B-cell lineage, *Ngp* and *S100a8* for neutrophils, and *Tpsb2* for mast cells (Figures S1C–S1E). Small clusters expressing markers of multiple lineages were designated as mixed cells, and proliferating cells were defined by *Mki67*, *Top2a*, and *Cdk1* expression. These populations were excluded from subsequent endothelial analyses. We next reclustered vascular cells to define endothelial and mural cell populations in greater detail (Figures 1C and S1F). Pericytes/vascular smooth muscle cells (VSMCs) were identified by *Pdgfrb*, *Cspg4*, *Tagln*, and *Acta2* expression, whereas lymphatic ECs were defined by *Prox1*, *Pdpn*, and *Lyve1* expression (Figures S1G and S1H). In adipose tissue, heart, and skeletal muscle, blood vascular ECs were classified into arterial, venous, and capillary ECs using established markers from previous endothelial atlases^11^: *Fbln5* and *Hey1* for arteries, *Vwf*, *Nr2f2*, and *Vcam1* for veins, and *Rgcc* and *Car4* for capillaries (Figures S1G and S1H). Liver ECs were annotated separately because of their distinct sinusoidal architecture. *Lyve1* and *Vwf* distinguished liver sinusoidal ECs from major-vessel ECs, and Vwf-positive ECs were further subdivided into central vein and portal vein ECs using *Rspo3*, *Bmp4*, *Adgrg6*, and *Nrg1* (Figures S1G and S1H).

**Figure 1.**
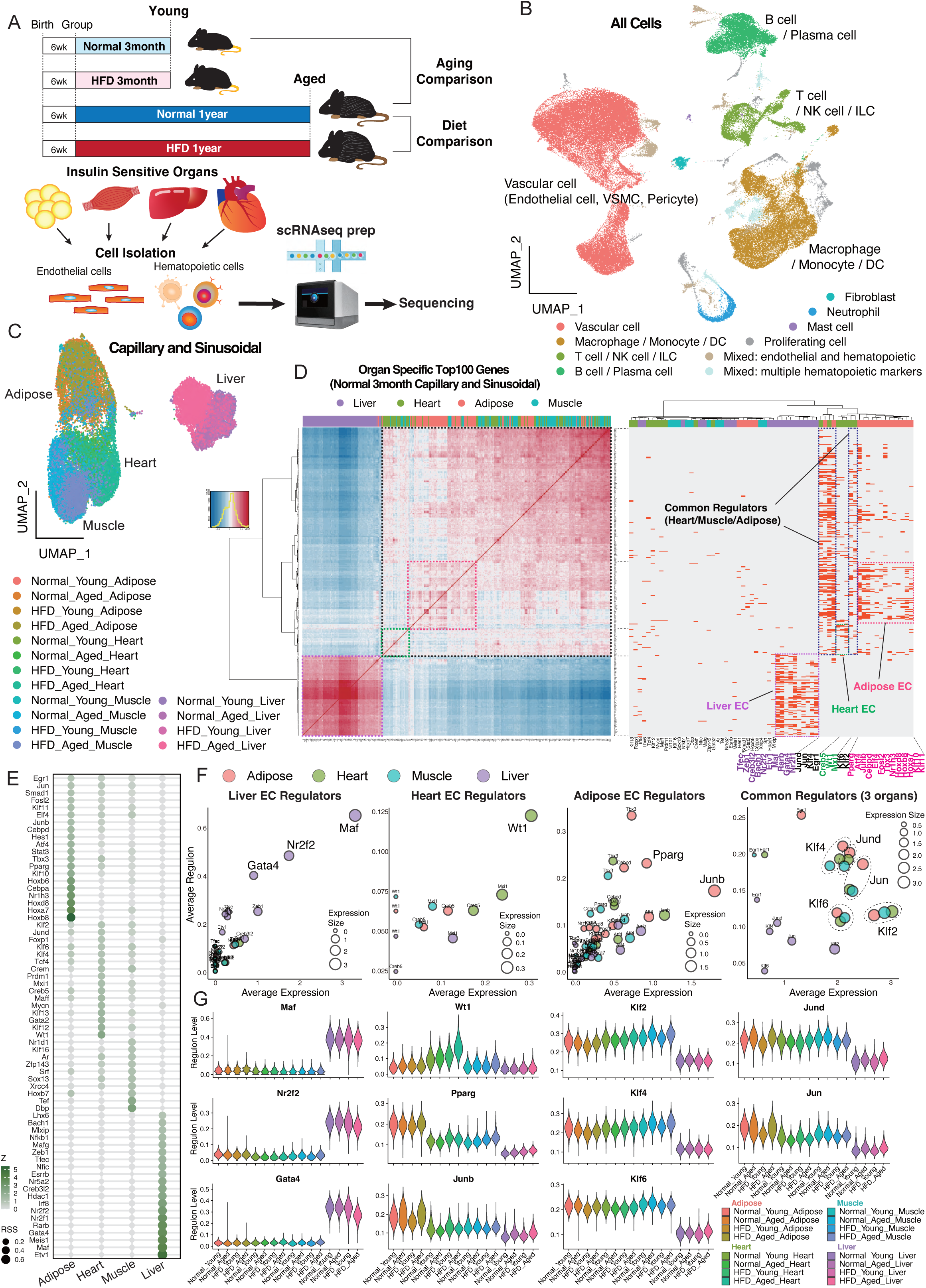
Single-cell profiling defines organ-specific endothelial identity across metabolic organs. (A) Experimental design. Male mice were fed normal diet or HFD and analyzed after 3 months or 1 year. Endothelial and hematopoietic cells from adipose tissue, skeletal muscle, liver, and heart were profiled by scRNA-seq. n = 1–2 mice per group. (B) UMAP of 141,590 quality-controlled cells. Major vascular, stromal, and hematopoietic populations are annotated; mixed-lineage clusters were excluded from downstream analysis. (C) Reclustering of capillary endothelial cells (ECs) from adipose tissue, skeletal muscle, and heart, and liver sinusoidal ECs. (D) Correlations of the top 100 organ-enriched endothelial genes in young normal-diet-fed mice. Spearman correlation and Ward.D2 clustering were used. (E) SCENIC regulon activity in young normal-diet-fed ECs. Regulons and cell types with RSS (Regulon Specificity Score) > 0.01 are shown. (F) The mean regulon AUC and RNA expression of representative organ-specific and shared regulons. Colors denote organs. (G) Violin plots of representative regulon AUC values across 16 organ-condition groups.

We first asked whether organ-specific endothelial identity was maintained under chronic stress. To focus on functional exchange vessels, we extracted capillary ECs from adipose tissue, skeletal muscle, and heart tissue, together with liver sinusoidal ECs, and reclustered them (Figure 1C). These cells clustered primarily by organ rather than by stress condition, indicating that organ-specific endothelial identity is largely preserved during aging and HFD loading. Consistent with these findings, organ-enriched endothelial genes exhibited strongly tissue-dependent expression patterns, with liver sinusoidal ECs forming the most distinct group and adipose, heart, and skeletal muscle ECs showing closer similarity with selected organ-biased modules (Figure 1D). To identify transcriptional regulators underlying these organ-specific endothelial states, we applied single-cell regulatory network inference and clustering (SCENIC)^18^ to samples of the four organs of young animals fed a normal diet (Figure 1E). We focused on transcription factors associated with organ-enriched endothelial genes or shared endothelial regulatory programs and compared both their inferred regulon activity and RNA expression across tissues. This parallel assessment helped support the interpretation of regulon-based organ-specificity (Figure S1I). This analysis revealed Maf, Nr2f2, and Gata4 as liver-enriched regulators; Wt1 as a heart-enriched regulator; Pparg and Junb as adipose-enriched regulators; and Klf2, Klf4, Klf6, Jund, and Jun as shared regulators among adipose tissue, heart, and skeletal muscle (Figures 1F and 1G). The enrichment of Wt1, Gata4, and Maf was consistent with previous reports on cardiac endothelial plasticity and hepatic sinusoidal endothelial specialization ^8,10,19,20^. These organ-associated regulatory patterns were maintained across stress conditions, although their relative activities varied with aging and HFD loading (Figure 1G). These results suggest that transcription factor programs underlying organ-specific endothelial identities are relatively resistant to aging- and diet-induced stress.

### Aging-mediated gene expression and transcriptional regulation in endothelial cells

To define aging-associated endothelial changes across organs, we reclustered capillary ECs from adipose tissue, skeletal muscle, and heart together with liver sinusoidal ECs from young and aged mice maintained on a normal diet (Figure 2A). For each organ, we selected the top 100 genes whose expression was upregulated in aged ECs (Figure S2A). Gene Ontology analysis of these organ-specific gene sets revealed similar functional enrichment across organs, with recurrent terms related to innate immune responses, inflammatory regulation, and interferon-associated pathways (Figure 2B). However, a comparison of the top 100 genes revealed that the individual genes were only partially shared among organs, and a substantial fraction remained organ-specific (Figure 2C). Thus, aging induced related functional programs across organs, but the responsible gene sets were not identical. We next asked which transcriptional regulators were associated with these aging-induced gene changes. SCENIC analysis was used to rank regulons activated in aged ECs from each organ (Figure 2D). Although several highly ranked regulons were shared across organs, others exhibited organ-biased patterns. To connect these candidate regulons with aging-induced genes, we integrated the top 100 aging-upregulated genes and the top 10 aging-associated regulons from each organ and calculated SCENIC-derived coexpression weights between the resulting union of genes and regulons. This analysis revealed four gene–regulon modules, designated Aging_1 to Aging_4, that captured strong relationships between organ-variable genes and their candidate upstream regulons (Figure 2E). The distribution of genes assigned to each module further supported the presence of both shared and organ-biased aging programs (Figure S2B). The Aging_1 module contained broadly induced genes and was associated with interferon-related regulons, including Irf7, Irf9, Stat1, and Stat2. The Aging_2 module was enriched for adipose-biased aging genes and was linked to Junb. The Aging_3 module contained genes preferentially altered in heart and skeletal muscle ECs and was associated with Klf family regulons, whereas the Aging_4 module exhibited a more restricted pattern involving muscle and liver ECs (Figure 2E). To validate whether these modules represented coherent single-cell transcriptional programs, we calculated signature scores for Aging_1 to Aging_4 using expression-matched control genes. These scores reproduced the organ- and age-dependent patterns inferred from the coexpression analysis, confirming that each Aging module reflected a distinct aging-associated endothelial state (Figure S2C). Representative genes from each module exhibited expression patterns consistent with the corresponding module scores (Figure S2D), and gene ontology analysis of the Aging_1–Aging_4 modules further supported their functional differences (Figure S2E).

**Figure 2.**
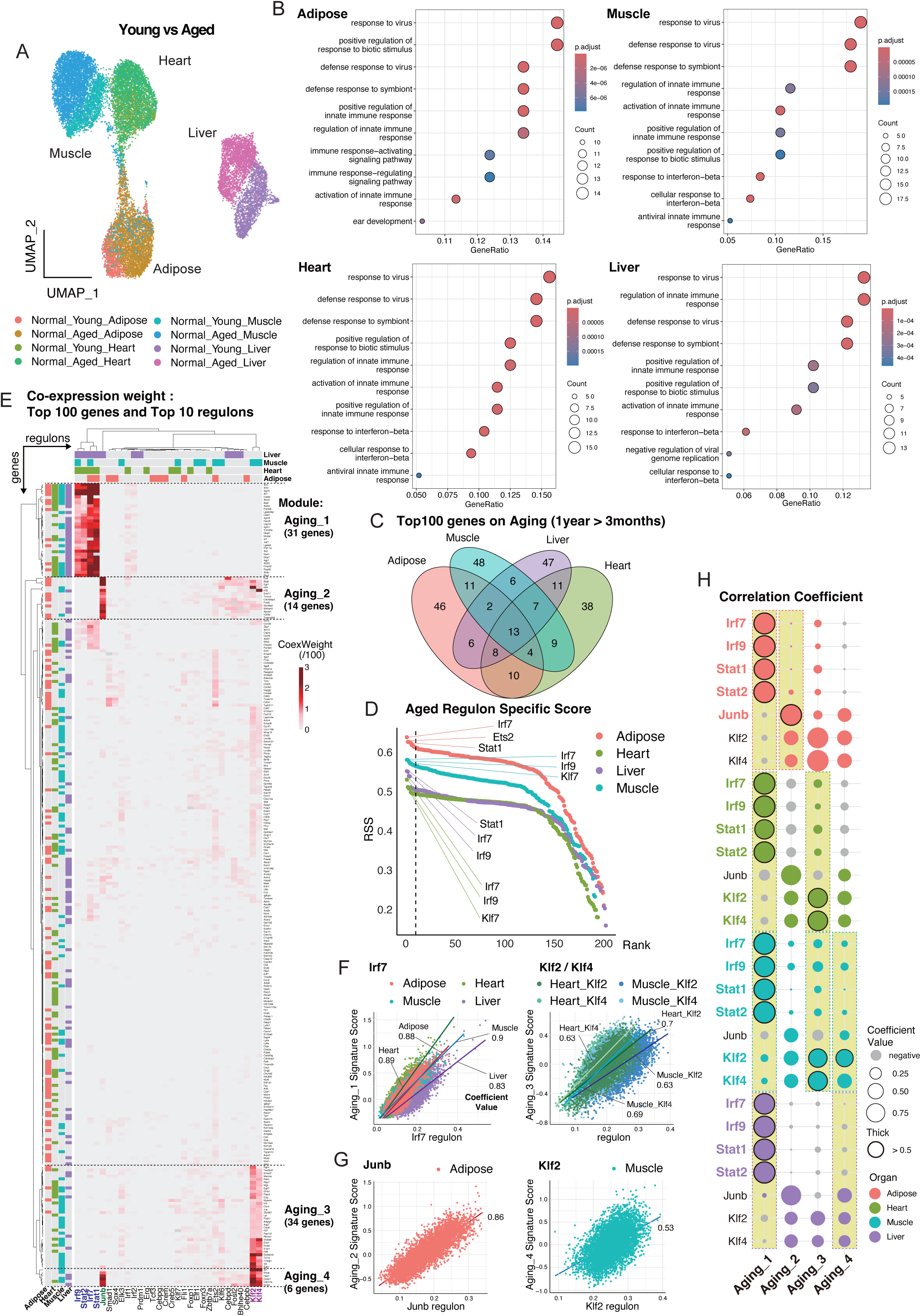
Organ-specific transcriptional networks associated with endothelial aging. (A) Reclustering of ECs from young and aged normal diet-fed mice. (B) Gene Ontology analysis of the top 100 aging-upregulated endothelial genes in each organ. (C) Overlap of the top 100 aging-upregulated genes across organs. (D) Organwise SCENIC ranking of aging-enriched regulons; the top regulons were merged for downstream analysis. (E) Coexpression weight matrix between the top aging-upregulated genes and aging-associated regulons, defining the Aging_1–Aging_4 modules. (F and G) Single-cell correlations between regulon activity and matched Aging module scores. Pearson correlation coefficients are indicated. (H) Summary matrix of Pearson correlations between regulon activity and Aging_1–Aging_4 scores. The circle size indicates the correlation coefficient. Yellow shading indicates organs contributing >30% of the genes in the corresponding module, as defined in Figure S2B.

Finally, we examined whether the candidate regulons were quantitatively linked to the corresponding aging module scores at single-cell resolution. For each organ, we correlated the aging module scores with the SCENIC-derived regulonAUC values, a per-cell measure of transcription factor regulon activity. Representative scatter plots revealed positive correlations between module scores and matched regulon activities, including between Aging_1 and Irf7, between Aging_2 and Junb, and between Aging_3/4 and Klf family regulons (Figures 2F, 2G, and S2F). A summary correlation matrix confirmed single-cell-level relationships between regulon activity and target gene module expression across Aging_1–Aging_4 and organs (Figure 2H). Notably, the Aging_1 module was enriched for interferon-stimulated genes (ISGs), including Ifit1, Ifit3, Isg15, Rsad2, Usp18, and Oasl2. Consistent with these findings, Irf7, Irf9, Stat1, and Stat2 emerged as regulators shared across organs and associated with this ISG-enriched aging module. Together, these findings indicate that endothelial aging is organized by both shared and organ-biased transcriptional regulatory modules.

### Diet-mediated gene expression and transcriptional regulation in endothelial cells

To define diet-induced endothelial changes across organs, we reclustered capillary ECs from adipose tissue, skeletal muscle, and heart tissue together with liver sinusoidal ECs from normal diet and 1-year HFD-fed mice (Figure 3A). Organwise analysis of HFD-upregulated endothelial genes revealed more limited overlap in gene ontology enrichment than in the aging response, indicating stronger organ-specificity under diet stress (Figures 3B and S3A). Related programs involving cell differentiation, oxidative stress, lipid metabolism, and tissue remodeling were nevertheless detected across several organs, whereas gene-level overlap remained low (Figure 3C). We next ranked the HFD-enriched regulons in each organ using SCENIC and found predominantly organ-biased regulatory patterns, with Pparg and Cebpd emerging as recurrent regulators across multiple tissues (Figure 3D). An integration of the HFD-upregulated genes with the diet-associated regulons revealed seven gene–regulon modules, Diet_1 to Diet_7, that linked diet-variable genes to candidate upstream regulators (Figure 3E). These modules showed distinct organ distributions, including multi-organ, adipose tissue, heart, skeletal muscle-enriched, heart-biased, and adipose-biased patterns (Figure S3B). Thus, HFD stress can engage partially shared transcription factors, such as Pparg and Cebpd, while redirecting their downstream targets to organ-specific endothelial programs.

**Figure 3.**
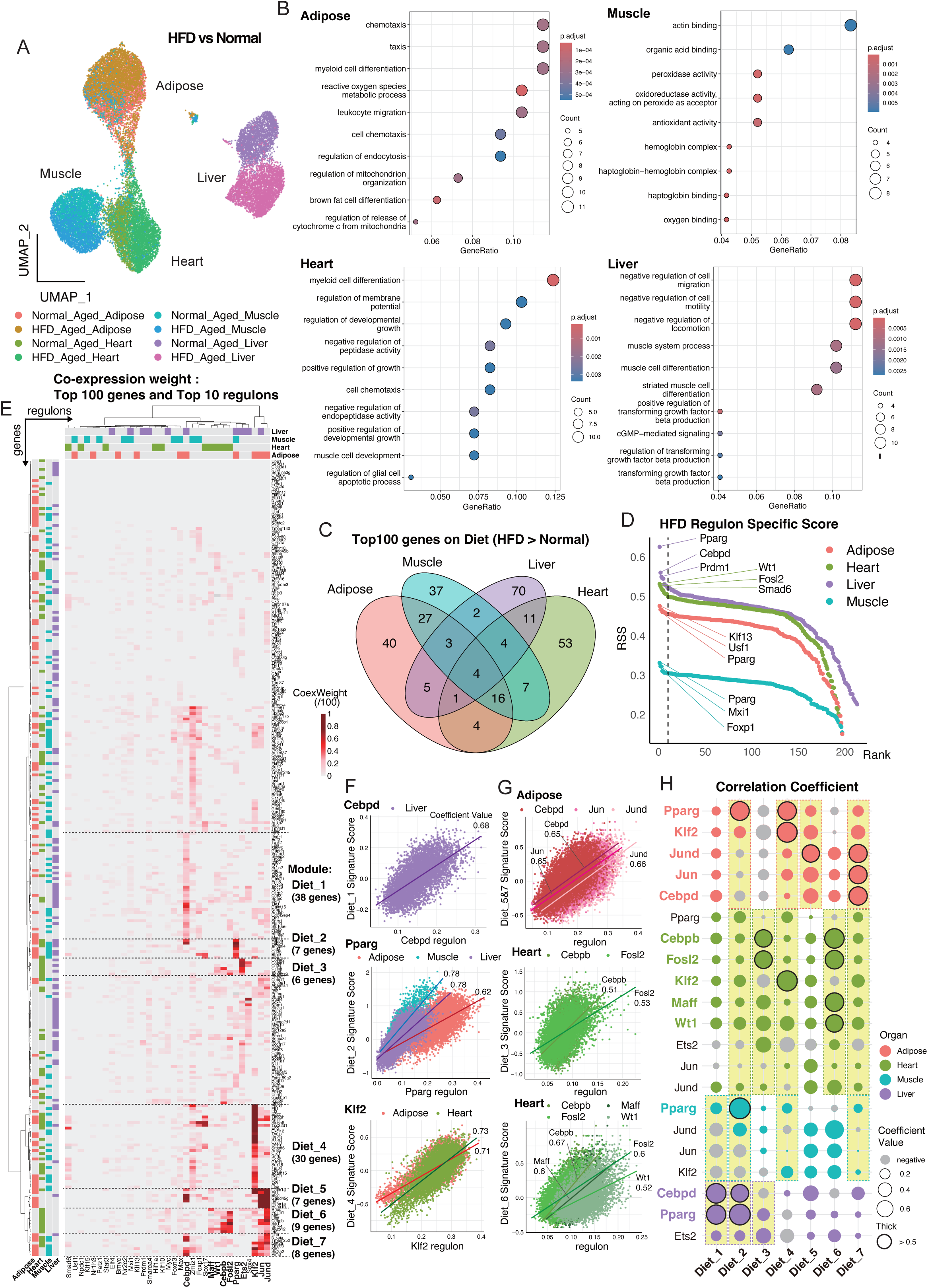
Organ-specific transcriptional networks associated with diet-induced metabolic stress. (A) Reclustering of ECs from 1-year-old normal-diet-fed and HFD-fed mice. (B) Gene Ontology analysis of the top 100 HFD-upregulated endothelial genes in each organ. (C) Overlap of the top 100 HFD-upregulated genes across organs. (D) Organwise SCENIC ranking of HFD-enriched regulons; the top regulons were merged for downstream analysis. (E) Coexpression weight matrix between the top HFD-upregulated genes and HFD-associated regulons, defining the Diet_1–Diet_7 modules. (F and G) Single-cell correlations between regulon activity and matched Diet module scores. Pearson correlation coefficients are indicated. (H) Summary matrix of Pearson correlations between regulon activity and Diet_1–Diet_7 scores. The circle size indicates the correlation coefficient. Yellow shading indicates organs contributing >30% of the genes in the corresponding module, as defined in Figure S3B.

We then calculated signature scores for Diet_1 to Diet_7, which reproduced the organ- and diet-dependent patterns inferred from the coexpression analysis and highlighted the stronger organ selectivity of the diet response (Figure S3C). Representative genes and gene ontology analysis supported the module assignments and functional diversity, including lipid metabolic processes in Diet_2 and angiogenesis- or morphogenesis-related programs in heart and muscle modules (Figures S3D and S3E). Correlation analysis further linked diet module scores to matched regulon activities at single-cell resolution, with representative examples involving Cebpd, Pparg, Jun/Jund, Klf2, Fosl2, Maff, and Wt1 depending on the organ and module (Figures 3F and 3G). A summary matrix confirmed that recurrent diet-associated regulons were linked to distinct diet modules across tissues (Figure 3H). Thus, HFD stress reshapes endothelial gene regulation in a more tissue-restricted manner than aging does, with partially shared upstream regulators driving divergent endothelial outputs across organs.

### Stress-specific and vascular-bed-shared regulon activation under aging and metabolic stress conditions

The preceding analyses revealed stress-associated endothelial regulons in aging and HFD comparisons. We then projected these regulons onto all 16 organ-condition groups using SCENIC-derived regulonAUC values to determine whether their activities reflected shared, organ specific, or stress-specific patterns (Figure 4A). Most regulons showed context-dependent activity rather than uniform activation across stresses. IFN-related regulons, including Irf7, Irf9, Stat1, and Stat2, were preferentially activated under aging conditions, whereas diet-associated regulons showed more organ-biased and stress-dependent patterns. Thus, endothelial transcriptional stress responses are shaped by both stress identity and tissue context. Among the aging-associated regulons, Irf7, Irf9, Stat1, and Stat2 showed the strongest positive correlations with one another at the single-cell level (Figure 4B), indicating coordinated activation of a type I IFN regulatory axis in aged ECs. Because STAT1, STAT2, and IRF9 mediate interferon-responsive transcription, whereas IRF7 contributes to type I interferon induction and amplification, these findings suggest that aged ECs engage both IFN-response and IFN-amplifying programs.

**Figure 4.**
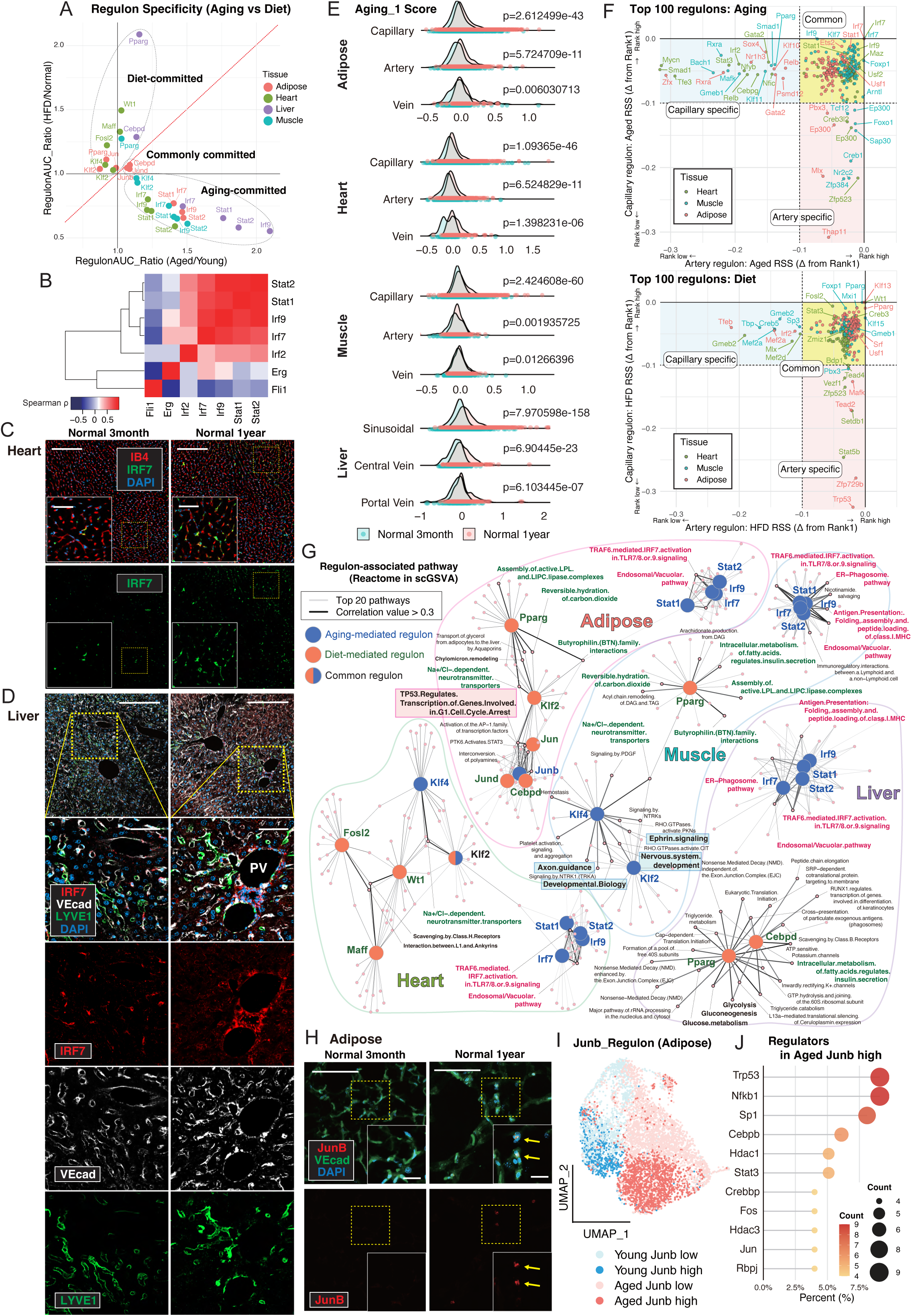
Organ-specific and vascular-bed-shared transcriptional regulon activation under aging and metabolic stress. (A) SCENIC-based comparison of representative aging- and diet-associated regulons across all organ-condition groups. (B) Single-cell correlation analysis of stress-associated regulon activities. Color indicates pairwise Spearman correlations of regulon activity. (C) Aging_1 signature scores across endothelial subtypes and age groups. (D) IRF7 immunofluorescence in aged tissues. PV, portal vein. Scale bars: low magnification, 200 µm; high magnification, 50 µm. (E) UMAP of arterial ECs from adipose tissue, heart tissue, and skeletal muscle. (F) Comparison of stress-responsive regulons between capillary and arterial ECs. Axis values indicate differences from the top-ranked stress-associated Δ value, with smaller differences indicating greater relative stress-responsiveness within each EC subset. (G) Network linking regulon activity to scGSVA-derived Reactome pathway scores. Edge thickness indicates Spearman correlation strength; red and green pathway labels indicate multi-organ aging and diet regulon links, respectively. (H) JunB immunofluorescence in adipose tissue. Scale bars: low magnification, 100 µm; high magnification, 50 µm. (I) Reclustering of adipose capillary ECs under normal diet conditions into Junb-high and Junb-low populations using a Junb regulon activity cutoff of 0.15, corresponding to the midpoint of the activity range shown in Figure S4J. (J) TRRUST upstream regulator analysis of genes enriched in aged Junb-high adipose ECs.

Immunostaining revealed that the number of IRF7-positive vascular ECs in heart and skeletal muscle increased with age (Figures 4C and S4A), and UMAP projection of Aging_1 scores similarly indicated that IFN activation was concentrated in a subset of ECs (Figure S4B). In the liver, IRF7-positive vascular and perivascular signals were also more prominent in aged tissue than in young tissue (Figure 4D). Consistent with these findings, the Aging_1 module scores increased with age across capillary, arterial, venous, and organ-specialized endothelial populations, indicating that the aging-associated IFN program extended beyond the capillary and sinusoidal endothelial populations used for initial module discovery (Figure 4E). Given the pathological importance of arterial endothelial remodeling, we further examined arterial ECs. Because liver arterial ECs were rarely recovered, this analysis was performed on adipose tissue, heart tissue, and skeletal muscle (Figure S4C). Using the same top 100 gene-based strategy, we found that arterial stress-responsive genes largely overlapped with the capillary endothelial programs, whereas artery-selective genes were limited and exhibited little cross-organ overlap (Figures S4D–S4G). SCENIC analysis further revealed that most stress-responsive regulons had similar relative behavior between arterial and capillary ECs (Figure 4F). These findings indicate that major endothelial stress regulons are broadly shared across vascular beds.

To link regulon activation to functional pathway states, we performed single-cell pathway enrichment analysis using scGSVA and correlated pathway scores with regulonAUC values. For each regulon, the top correlated Reactome pathways were extracted, and highly correlated regulon–pathway relationships were summarized as a network (Figures 4G and S4H). Aging-associated Irf/Stat regulons were linked to immune-related pathways, including antigen processing and presentation, across multiple organs. This pattern was consistent with the gene ontology results for the Aging_1 module, which exhibited enrichment of immune and interferon-related programs (Figure S2E). In contrast, diet-associated regulons exhibited greater organ-specificity. Consistent with the results of the gene–regulon module analysis shown in Figure 3, this single-cell pathway analysis further resolved how tissue-biased regulon activity is linked to distinct functional pathway states under metabolic stress. Pparg-associated activity was linked to lipid handling and metabolic pathways, whereas Klf and AP-1 family regulons were associated with more tissue-selective pathway relationships. Among these pathway relationships, p53-related cell cycle arrest and stress response pathways were selectively evident in adipose ECs and were associated with coordinated AP-1 regulon activity (Figure 4G). This pattern is notable because our previous studies revealed that endothelial p53 signaling contributes to vascular dysfunction under hyperglycemic, ischemic, and obesity-related stress conditions^21,22^, suggesting that even canonical endothelial stress pathways can be preferentially engaged in a tissue-restricted manner. Consistently, Junb regulon activity increased with age and was further increased under HFD conditions (Figure S4I), and immunostaining confirmed an increased nuclear JunB signal in aged adipose ECs (Figure 4H). In fact, in the aging comparison, upstream regulator analysis of the aged Junb-high fraction positioned Trp53 as the top regulator relative to the Junb-low fraction (Figures 4I, 4J, and S4J). Together, these findings indicate that endothelial stress remodeling involves both broadly shared aging programs, represented by coordinated IFN regulon activation, and organ-biased stress states, including a distinct adipose endothelial fraction marked by p53 associated activation.

### Cell–cell interaction analysis highlights stress-specific endothelial–immune communication

To determine whether endothelial stress programs were associated with changes in surrounding immune cells, we performed single-cell-based cell–cell interaction (CCI) analysis using CellChat^23^ between vascular endothelial subsets and the Macrophage, T-cell, and B-cell lineages (Figure 5A). Stress-responsive signaling pathways were extracted by comparing aging and HFD conditions with their corresponding controls (Figure S5A). When altered signaling pathways were classified by sender and receiver directionality, endothelial-to-endothelial communication accounted for a large fraction of the stress-responsive interactions (Figures 5B and S5B). Among capillary-to-capillary signals, the EPHB pathway was increased in skeletal muscle ECs under both aging and HFD stress conditions (Figure S5C). EPHB-associated ligands and receptors, including Efnb1, Efnb2, Epha4, and Ephb4, were linked to ephrin signaling and Klf2/Klf4-associated regulon activity in aged skeletal muscle ECs (Figure S5D). Building on prior links among ephrin–EphB signaling, endothelial network formation, and communication-driven transcriptional response^23–26^, these findings identify ephrin–EphB signaling as a stress-responsive endothelial communication axis aligned with organ-biased regulatory programs.

**Figure 5.**
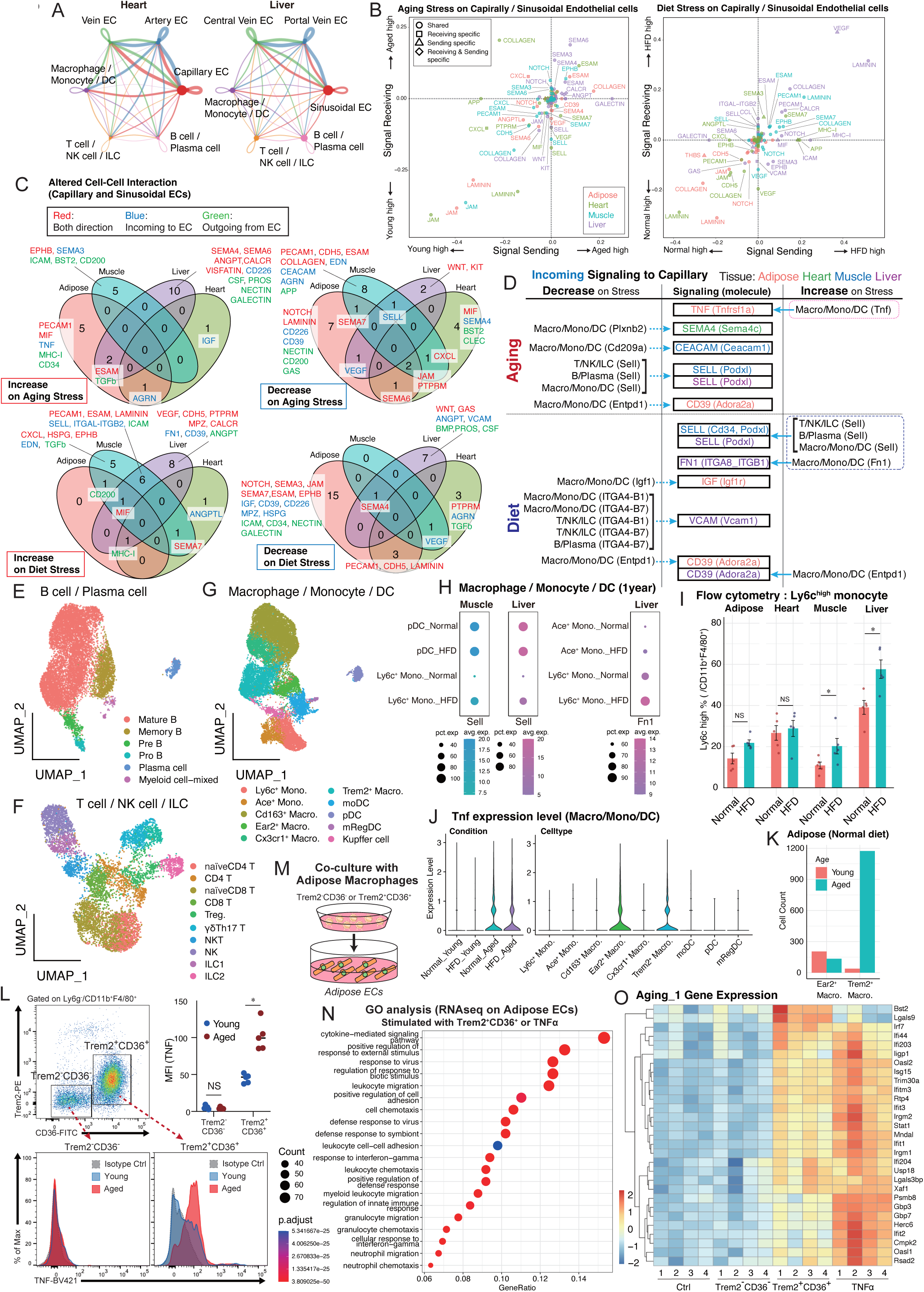
Cell–cell interaction analysis highlights stress-specific endothelial–immune communication. (A) Cell–cell interaction networks between endothelial subsets and immune lineages. (B) Differentially regulated signaling pathways in capillary and sinusoidal ECs under aging or HFD stress. (C) Summary of altered CCI pathways classified as capillary-to-capillary, endothelial outgoing, or endothelial incoming. (D) Incoming immune-to-endothelial signals altered under aging or HFD stress. The central boxes indicate signaling pathways and corresponding endothelial receiving molecules, whereas the outer labels indicate inferred hematopoietic sender populations and their input molecules in parentheses. (E–G) UMAP subclustering of the Macrophage, T-cell, and B-cell lineages. Treg.: regulatory T, Mono.: monocyte, Macro: macrophage, DC: dendric cell. (H) Sell and Fn1-associated source signals in immune subsets under HFD stress. (I) Flow-cytometric quantification of Ly6c-high monocytes under normal diet and HFD conditions. Cells were gated from Ly6G^−^CD11b^+^F4/80^+^ populations; n = 5 mice per group. (J) Tnf expression in adipose Macrophage lineage subsets across conditions. (K) Ear2^+^ and Trem2^+^ macrophage numbers in young and aged adipose tissue. (L) TNF protein expression in Trem2^−^CD36^−^ and Trem2^+^CD36^+^ adipose macrophages. Cells were gated from Ly6G^−^CD11b^+^F4/80^+^ populations; n = 5 mice per group. (M) Adipose ECs stimulation design using macrophage co-culture or recombinant TNFα. (N) Gene Ontology analysis of genes with upregulated expression after Trem2^+^CD36^+^ macrophage co-culture or TNFα stimulation. (O) Aging_1 gene expression after macrophage co-culture or TNFα stimulation.

At the pathway level, endothelial incoming and outgoing signals showed limited overlap across organs, indicating that stress-responsive vascular communication is largely organ- and context-dependent (Figure 5C), in line with our recent finding that tissue context shapes organ-specific immune cell programs^27^. We next focused on incoming signals from immune cells to capillary or sinusoidal ECs and identified stress-induced alterations in immune-to-endothelial communication pathways under aging or HFD conditions (Figure 5D). To interpret these immune-to-endothelial communication patterns at higher cellular resolution, we subclustered the B-cell, T-cell, and Macrophage lineages (Figures 5E–5G and S5E–S5J). Although Sell was detectable in several T-cell and B-cell subsets, these populations exhibited limited diet-associated changes in skeletal muscle (Figures S5K and S5L). In contrast, Ly6c^+^ monocytes exhibited increased Sell and Fn1-associated signals in skeletal muscle and liver under HFD conditions (Figure 5H). Consistent with these findings, single-cell analysis and flow cytometry confirmed increased Ly6c^+^ monocyte accumulation in these organs after HFD loading (Figures 5I, S5M, and S5N), in line with previous reports of CCR2-dependent Ly6C^+^ monocyte recruitment in obesity and steatohepatitis-associated inflammation^28–30^.

We then examined the aging-associated TNF signal in adipose tissue. *Tnf* expression in the Macrophage lineage increased with age regardless of diet condition (Figure 5J). Subcluster analysis revealed that Trem2^+^ macrophages were a major macrophage population associated with this age-dependent increase, and that Trem2^+^ macrophage numbers markedly increased in aged adipose tissue under normal diet conditions (Figure 5K). Trem2^+^ macrophages, also referred to as lipid-associated macrophages (LAMs), play a crucial role in lipid metabolism and metabolic tissue remodeling^31^. On the basis of previous reports using CD36 as a marker of lipid-associated macrophage states, we used Trem2 and CD36 to enrich this population for flow cytometric analysis. Flow cytometry confirmed that TNF protein expression increased with aging specifically in Trem2^+^CD36^+^ macrophages, whereas Trem2^−^CD36^−^ macrophages showed little change (Figure 5L).

To test whether aging-associated adipose macrophages could induce endothelial stress programs, we cocultured adipose ECs with Trem2^+^CD36^+^ or Trem2^−^CD36^−^ adipose macrophages isolated from aged mice or stimulated them with recombinant TNFα (Figure 5M). Bulk RNA-seq revealed that Trem2^+^CD36^+^ macrophages and TNFα induced overlapping transcriptional changes, whereas Trem2^−^CD36^−^ macrophages had a weaker effect (Figures S5O and S5P). Gene Ontology analysis of genes induced by Trem2^+^CD36^+^ macrophages or TNFα revealed enrichment of cytokine-mediated signaling, response to external stimulus, response to virus, and leukocyte migration-related terms (Figure 5N), which was consistent with the inflammatory and interferon-associated enrichment observed in aging-responsive endothelial genes (Figure S2A). We then examined individual Aging_1 module genes defined in Figure 2E. These genes were broadly induced by Trem2^+^CD36^+^ macrophages and TNFα, but not by Trem2^−^CD36^−^ macrophages (Figure 5O). Thus, Trem2^+^CD36^+^ adipose macrophages provide a TNF-rich immune input capable of activating the endothelial Aging_1 program. Although TNFα is not a canonical type I interferon ligand, previous work has shown that TNFα can converge on interferon-stimulated gene transcription and cooperate with interferon signaling^32^, supporting a role for macrophage-derived TNF in endothelial IFN-related amplification.

### Spatial transcriptomics maps perivascular ISG clusters in aged tissues

Single-cell and CCI analyses revealed organ-biased endothelial stress responses, a shared aging-associated IFN program, and immune-derived TNF input linked to Aging_1 activation in ECs. We next performed spatial transcriptomic analysis in young and 24-month-old very aged mice to determine how this program is organized within heart and adipose tissue architecture. Spatial cell identities were assigned by label transfer using the single-cell datasets generated in this study and mouse aging atlas resources, including Tabula Muris Senis and our previously reported aging-related dataset^16,33^. Across two biological replicates per tissue and age group, we annotated major cell populations in both heart and adipose tissue (Figures 6A, 6B, S6A, and S6B). Projection of the Aging_1 module score onto spatial maps revealed age-associated increases in both heart and adipose tissue (Figure S6C). In the heart, high Aging_1 scores appeared as spatially restricted foci, not as diffuse tissue-wide signals (Figure 6C), and similar focal enrichment was observed in adipose tissue (Figure 6D). Targeted in situ gene expression profiling further confirmed age-associated focal accumulation of Aging_1-high cells across multiple organs, with prominent clusters in the heart (Figure S6D). We refer to these focal domains, including Aging_1-high regions and their local surroundings, “ISG clusters,” reflecting enrichment of interferon-stimulated genes within the Aging_1 module.

**Figure 6.**
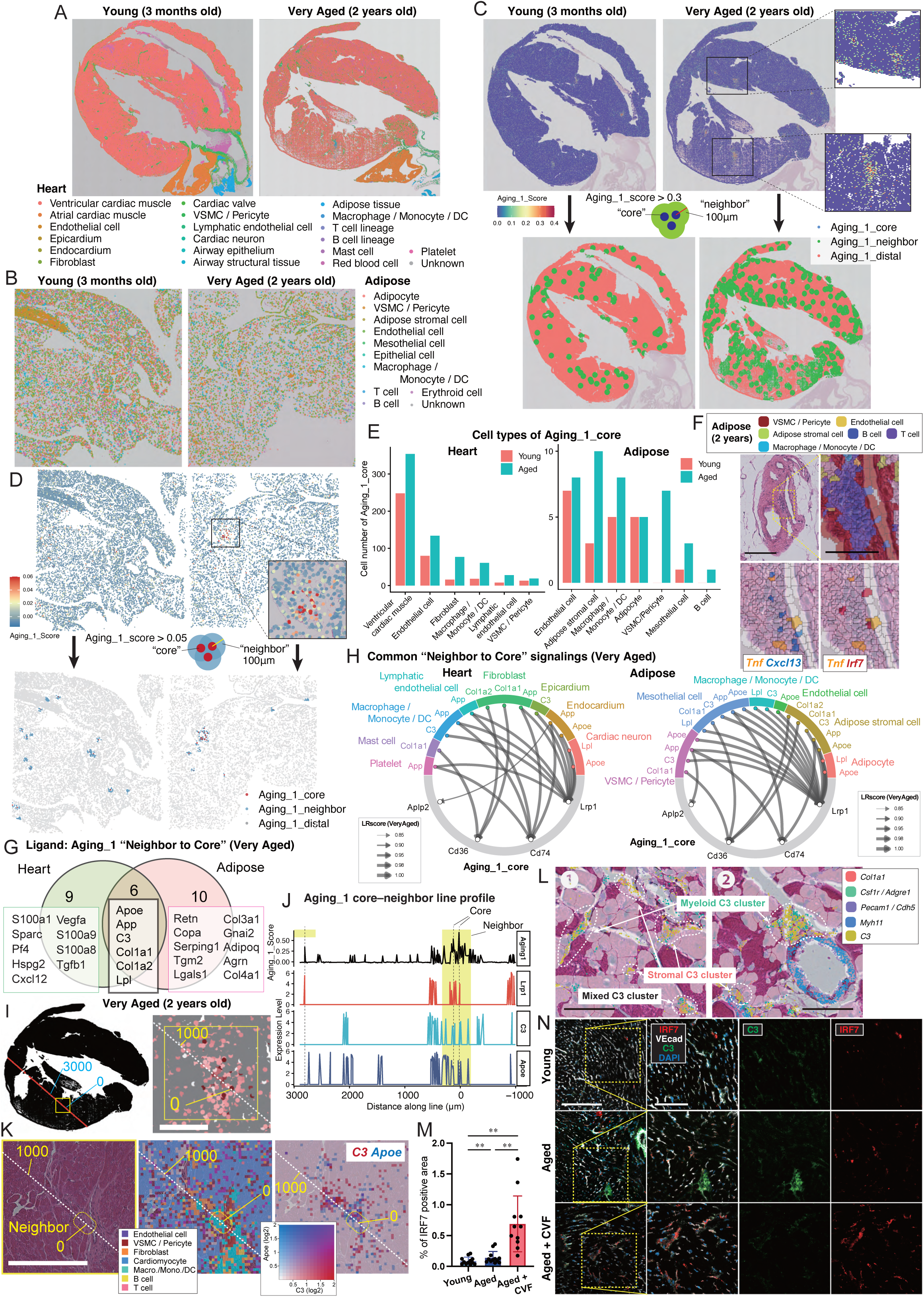
Spatial transcriptomics maps perivascular ISG clusters in aged tissues. (A) Spatial annotation of young and very aged heart sections using Visium HD 8 µm bin data. (B) Spatial annotation of young and very aged adipose tissue using Visium HD cell segmentation data. (C) Aging_1 scores in the ventricular region. Aging_1 core, neighbor, and distal cells were defined by high scores, ≤100 µm from the core, and remaining cells, respectively. (D) Aging_1 scores in adipose tissue. The compartments are defined as in (C). (E) Cell-type composition of Aging_1 core cells in heart and adipose tissue. (F) Spatial localization of Tnf, Cxcl13, and Irf7 in very aged adipose tissue. Scale bars: H&E overview, 100 µm; magnified regions, 50 µm. (G) Ligands enriched in Aging_1 neighbor-to-core signaling in very aged heart and adipose tissue. (H) Common Aging_1 neighbor-to-core ligand–receptor signals. Edge thickness indicates the LIANA-derived LR score. (I) Spatial line across an Aging_1 core–neighbor region in very aged heart. One high Aging_1 core was set as 0 µm, and the numbers indicate the distance from this point. Scale bar, 0.5 mm. (J) Line-profile analysis of bins along the spatial line in (I), showing the Aging_1 score and gene expression levels. The neighboring region is highlighted in yellow. (K) Magnified view of the yellow-boxed region in (I). Left, H&E; middle, cell-type annotation; right, C3 and Apoe expression. Scale bar, 1 mm. (L) Magnified Xenium views of regions ❶ and ❷ indicated in Figure S6K. C3-expressing clusters contained Col1a1-expressing stromal cells, Csf1r/Adgre1-expressing myeloid cells, or mixed stromal–myeloid components. Scale bar, 50 µm. (M) Quantification of IRF7-positive ECs in cardiac sections from aged mice treated with PBS or CVF. Three regions were analyzed per mouse, with four mice per group, yielding 12 regions per group. P values were calculated using region-level measurements. **p < 0.001. (N) IRF7, VE-cadherin, C3, and DAPI immunofluorescence in young, aged, and C3-depleted aged hearts. Scale bars: overview, 200 µm; magnified regions, 100 µm.

To define the organization of ISG clusters, Aging_1-high spatial units were designated as Aging_1_core, spatial units within 100 μm as Aging_1_neighbor, and the remaining units as Aging_1_distal (Figures 6C and 6D). We then examined the cellular composition of the Aging_1_core and Aging_1_neighbor compartments. In the heart, Aging_1_core included ventricular cardiac muscle, reflecting the surrounding myocardial architecture; among non-cardiomyocyte populations, ECs were most abundant, followed by fibroblasts and cells of the Macrophage lineage (Figure 6E). In adipose tissue, ECs, adipose stromal cells, cells of the Macrophage lineage, and VSMCs/pericytes contributed to Aging_1_core, and these populations increased with aging (Figure 6E). Aging_1_neighbor also expanded with aging and contained endothelial, fibroblast/stromal, VSMC/pericyte, and Macrophage lineage populations (Figures S6E and S6F). Together, these spatial and cellular features indicate that ISG clusters are frequently associated with vascular-associated compartments.

To localize the immune-derived TNF signal identified above, we examined aged adipose tissue. Tnf was detected in perivascular LAMs (Figure S6G). Notably, Tnf also accumulated in B cell-rich aggregates near arterial vascular structures (Figure 6F). These aggregates were accompanied by Cxcl13-expressing cells, consistent with features of TLS-like inflammatory organization in chronic inflammation^34,35^. *Irf7*-positive cells were also observed around *Tnf*-enriched regions, supporting a spatial association between local TNF-rich inflammatory input and endothelial IFN activation.

We next performed spatial CCI analysis using LIANA^36^ to map candidate ligand–receptor interactions directed toward Aging_1_core within ISG clusters. For each tissue, we extracted pathways that were stronger from Aging_1_neighbor to Aging_1_core than from Aging_1_distal to Aging_1_core (Figures S6H and S6I) tissue and focused on ligands increased in very aged tissues. Six ligands were shared between heart and adipose tissue: *C3*, *Apoe*, *App*, *Lpl*, *Col1a1*, and *Col1a2* (Figure 6G). Ligand–receptor visualization revealed that these shared incoming signals converged on several receptors expressed in Aging_1_core, including *Lrp1*, *Cd74*, *Cd36*, and *Aplp2* (Figure 6H). Among these receptors, *Lrp1* was the most recurrent Aging_1_core-side receptor across the shared ligand interactions in both organs, including interactions involving *C3* and *Apoe*. Analysis of these two *Lrp1*-linked ligands showed that *Apoe* in the Aging_1_neighbor compartment was enriched mainly in the Macrophage lineage, whereas C3 was prominent in fibroblast or stromal populations and was also detected in the Macrophage lineage (Figure S6J). We therefore focused on the spatial relationship between Lrp1-expressing Aging_1_core regions and C3- or Apoe-enriched Aging_1_neighbor compartments. Line-profile analysis across representative ISG clusters revealed that Lrp1 expression aligned with Aging_1_core regions in the very aged heart, whereas *C3* and *Apoe* were enriched in the surrounding Aging_1_neighbor compartment (Figures 6I and 6J). A similar pattern was observed in very aged adipose tissue (Figure S6K). A magnified view of the heart ISG cluster further revealed Aging_1_core regions positioned near spatially adjacent but largely distinct *C3-* and *Apoe-*expressing areas (Figure 6K). Targeted in situ gene expression profiling further showed that *C3*-expressing regions in the aged heart were organized as perivascular clusters that included *Col1a1*-expressing stromal cells, *Csf1r/Adgre1*-expressing myeloid cells, or mixed stromal–myeloid components (Figures 6L and S6L). Because *C3* may engage distinct receptor contexts, we examined the spatial relationships among *C3*, *Lrp1*, and *C3ar1*. Bivariate mapping revealed *C3-*expressing regions near Lrp1-positive areas, whereas C3ar1 exhibited a distinct distribution (Figure S6M). This finding suggested that C3 may be associated with Lrp1-linked scavenging or buffering responses, prompting us to test whether C3 activity modulates perivascular IFN inflammation. We therefore depleted C3 with cobra venom factor (CVF) and assessed cardiac IRF7 expression. IRF7-positive vascular regions increased with age, and C3 depletion further increased the number of IRF7-positive areas (Figure 6M and 6N). These findings indicate that aging-associated endothelial IFN activation is spatially organized into perivascular ISG clusters, where C3-associated signals may contribute to the local restraint of vascular IFN activation.

### Endothelial BST2 downstream of IFN signaling promotes anti-inflammatory macrophage differentiation

To connect stress-activated transcription factors with downstream effector molecules involved in cell–cell communication, we returned to the endothelial–immune CCI framework and examined CCI-associated molecules within stress-responsive regulons (Figure 7A). For each regulon–molecule pair, regulonAUC and target gene expression were correlated at the single-cell level and ranked by R². Among the highest-ranked pairs, IFN-related regulons, including Irf7, Irf9, Stat1, and Stat2, were linked to *Bst2* expression. This relationship was selectively evident in heart and skeletal muscle ECs (Figures 7B and 7C). *Bst2* expression increased with age in multiple endothelial cell subtypes, including capillary, arterial, and venous ECs, in these two organs (Figure 7D). BST2, also known as tetherin, is an interferon-inducible membrane protein originally characterized as an antiviral restriction factor that prevents the release of enveloped virions from infected cells^37–39^. More recent work has also implicated BST2–PIRA2 communication in vascular smooth muscle cell plasticity in the tumor microenvironment^40^. However, the functional meaning of BST2 induction in vascular ECs during aging-associated stress remains unclear. Other aging-associated endothelial molecules, including *Cxcl12* and *Lgals9*, were also identified (Figure S7A). In the heart, Cxcl12 increased with aging whereas *Ackr3*, also known as Cxcr7, decreased, suggesting that Cxcl12-related communication may extend beyond endothelial-to-endothelial signaling.

**Figure 7.**
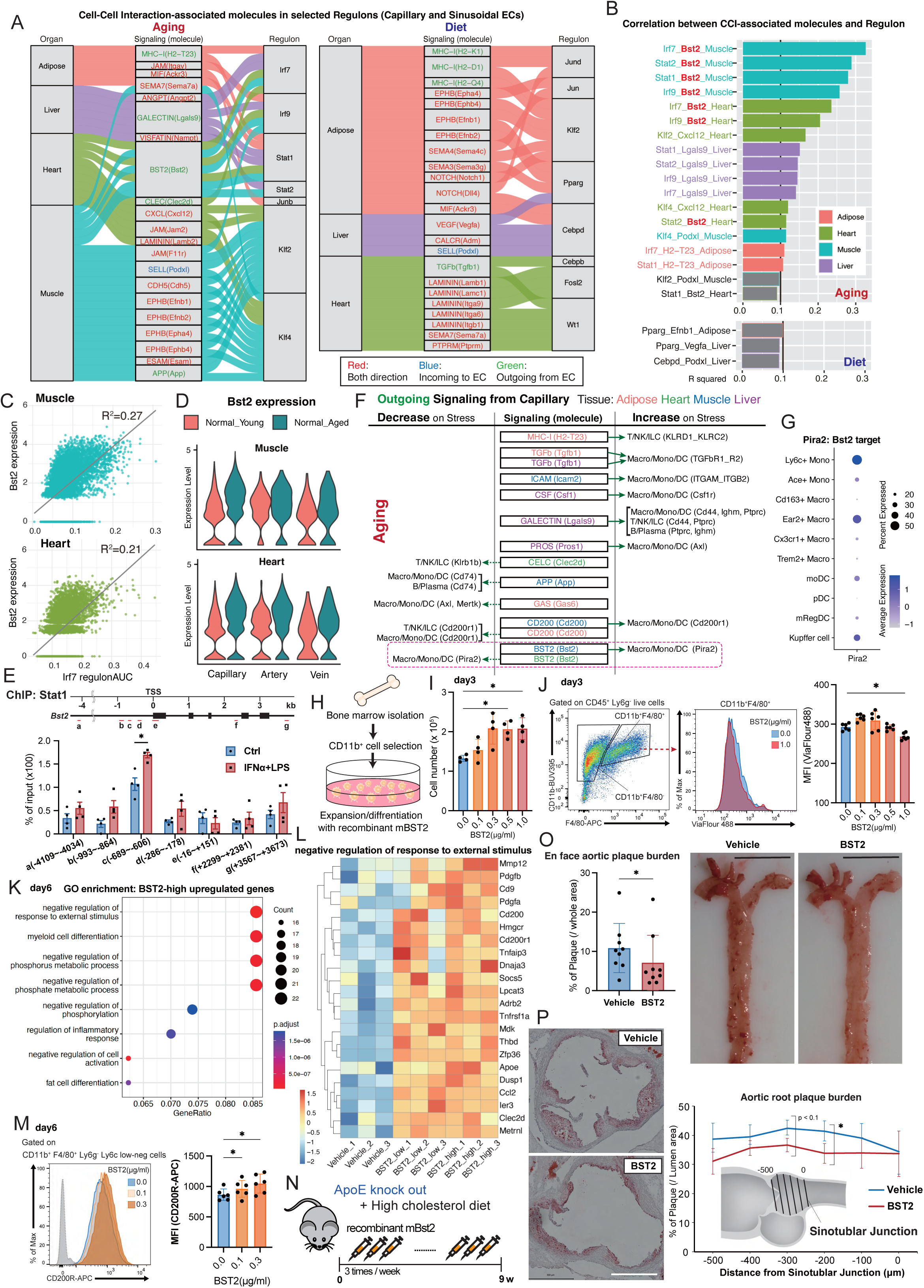
Endothelial BST2 downstream of IFN signaling promotes anti-inflammatory macrophage differentiation and suppresses atherosclerosis. (A) Sankey plot linking organs to CCI-associated effector pathways and molecules. Colors indicate endothelial outgoing, endothelial incoming, or bidirectional pathways. (B) Ranked regulon–effector molecule–organ combinations from correlation analysis. (C) Representative single-cell correlation between IFN-related regulon activity and *Bst2* expression in heart and skeletal muscle ECs. (D) *Bst2* expression in capillary, arterial, and venous ECs from heart and skeletal muscle. (E) STAT1 ChIP analysis at the *Bst2* locus in primary cardiac ECs treated with IFNα followed by LPS. (F) Outgoing endothelial-to-immune signaling under aging stress. The central boxes indicate signaling pathways and corresponding endothelial output molecules, whereas the outer labels indicate inferred immune recipient populations and their receiving molecules in parentheses. (G) *Pira2* expression across Macrophage lineage subsets. Dot size indicates the percentage of expressing cells; color indicates average expression. (H) BMDM differentiation assay using CD11b^+^ bone marrow cells and recombinant mouse BST2. (I) Total cell number on day 3 after recombinant BST2 stimulation. (J) ViaFluor 488 proliferation analysis of CD11b^+^F4/80^+^ and CD11b^+^F4/80^−^ cells. *p < 0.05. (K) Gene Ontology analysis of genes with upregulated expression in BST2-high cells on day 6. BST2-low, 0.1 µg ml^−1^; BST2-high, 1.0 µg ml^−1^ recombinant mouse BST2. (L) Genes contributing to the “negative regulation of response to external stimulus” term, highlighting Cd200r1. (M) CD200R mean fluorescence intensity in CD11b^+^F4/80^+^Ly6G^−^Ly6C^low-negative^ cells on day 6. *p < 0.05. (N) Recombinant BST2 administration regimen in high-cholesterol diet-fed ApoE-deficient mice. Recombinant BST2 was injected three times per week for 9 weeks. (O) En face Oil Red O analysis of the whole-aorta plaque burden in mice treated as in (N). n = 9 mice per group, 6 males and 3 females, pooled from three independent treatment cohorts. Scale bar, 5 mm. *p < 0.05. (P) Aortic root plaque analysis in mice treated as in (N). Representative sections at 200 µm from the sinotubular junction and serial-section quantification are shown. n = 5 mice per group, 2 males and 3 females, pooled from two independent treatment cohorts. Scale bar, 500 µm. *p < 0.05, two-sided t test.

We examined whether *Bst2* induction was directly linked to transcriptional regulation at the *Bst2* locus. *Bst2* was included among the Aging_1 genes, and STAT1 ChIP in primary cardiac ECs revealed increased binding upstream of the Bst2 transcription start site after IFNα and LPS stimulation (Figure 7E), supporting direct regulation by IFN-related signaling. We next analyzed stress-regulated outgoing endothelial signals to immune lineages. Bst2 was linked to signaling toward cells of the Macrophage lineage through Pira2, particularly in heart and skeletal muscle (Figures 7F and S7B). *Pira2* expression was highest in Ly6c^+^ monocytes (Figure 7G), suggesting that these cells are potential recipients, although the functional direction of this interaction could not be resolved by CCI analysis alone. Given previous reports linking BST2–PIRA2-related communication to immune regulation^40,41^, we tested the functional effect of BST2 using a BMDM differentiation assay. CD11b^+^ bone marrow cells were cultured under macrophage differentiation conditions with recombinant mouse BST2 (Figure 7H). BST2 increased total cell numbers on day 3 (Figure 7I) and increased the proliferation of CD11b^+^F4/80^+^ macrophage-lineage cells, whereas CD11b^+^F4/80^+^ cells were not significantly affected (Figures 7J and S7C). We subsequently examined transcriptional and phenotypic changes on day 6. Bulk RNA-seq revealed dose-dependent transcriptional responses to BST2 (Figure S7D), with enrichment of regulatory programs including negative regulation of response to external stimulus, myeloid cell differentiation, and negative regulation of inflammatory response (Figure 7K). *Cd200r1* was increased among the genes contributing to this regulatory signature (Figure 7L), which was consistent with increased CD200-related endothelial-to-Macrophage lineage signaling under aging stress in skeletal muscle (Figure 7F). Flow cytometry confirmed increased CD200R protein expression on a per-cell basis, whereas the fraction of CD200R-positive cells varied across samples (Figures 7M and S7F). Other macrophage markers, including CX3CR1, I-A/I-E, and CD206, were not substantially altered (Figure S7E and S7G). Thus, although BST2 is induced downstream of IFN signaling, it may contribute to a compensatory restraint mechanism by enhancing CD200R-associated regulatory features in macrophages.

The Bst2-associated communication route was identified in heart and skeletal muscle, two organs closely related to atherosclerotic vascular disease. In addition, the CD200–CD200R checkpoint has been reported to limit monocyte–macrophage activation and recruitment during atherosclerosis progression^42^. Given that BST2 increased CD200R expression during macrophage differentiation, we tested whether BST2 could influence atherogenesis *in vivo*. Recombinant mouse BST2 was administered to ApoE-deficient mice fed a high-cholesterol diet for 9 weeks (Figure 7N). Continuous BST2 treatment significantly reduced atherosclerotic plaque burden in the whole aorta and at the aortic root (Figures 7O, 7P, and S7H). We confirmed CD200R expression in aortic root lesional macrophages in both groups (Figure S7I). Together, these findings indicate that the aging-associated endothelial IFN program is not solely linked to inflammatory activation, but can also induce regulatory effector molecules. BST2 represents one such IFN-induced endothelial effector, linking inflammatory IFN activation to CD200R-associated macrophage regulation and suppression of atherosclerotic lesion formation.

## Discussion

In this study, we defined how vascular ECs integrate aging and metabolic overload across organs. By profiling endothelial and hematopoietic populations in adipose tissue, skeletal muscle, liver, and heart, we linked endothelial transcriptional remodeling to immune cell composition and cell–cell communication. Spatial transcriptomics further resolved vascular niches in which endothelial, stromal, and myeloid cells are organized around aging-associated inflammatory programs. These analyses indicated that vascular inflammaging is locally organized, with inflammatory activation and restraining signals coexisting within perivascular neighborhoods. Despite chronic systemic stress, organ-specific endothelial transcriptional features were relatively preserved. This finding is consistent with prior work showing that ECs are specialized by organ and vascular bed, and do not form a uniform vascular lining^6,11,14^, and that vascular bed identity is shaped by developmental and postnatal endothelial patterning programs^2,3,25,43,44^. Our data extend this view by showing that aging and metabolic overload affect preexisting vascular bed specialization to generate shared and organ-biased endothelial stress responses.

Aging and high-fat diet stress remodeled endothelial regulatory networks in distinct ways. Aging induced a broadly conserved interferon-related program involving Irf7, Irf9, Stat1, and Stat2, whereas high-fat diet stress preferentially engaged organ-biased lipid metabolic and remodeling programs, including Pparg and Cebpd-associated regulons. Consistent with this aging-related component, recent multi-organ endothelial single-cell analyses have similarly revealed that aging induces shared interferon-responsive and antigen-presentation programs across organs, including organs that overlap with those analyzed here^45^. Previous studies of endothelial aging and metabolic stress have also described the activation of inflammatory and interferon-responsive programs, together with the attenuation of angiogenic programs and endothelial regenerative capacity^17,46,47^. Our study extends these observations by showing that the endothelial IFN program is coupled to organ-specific endothelial–immune communication and is restricted to local vascular niches.

Although aging is often linked conceptually to cellular senescence, chronological aging does not correspond to a uniform senescent cell state. p53 signaling illustrates this distinction, as previous studies, including work from our groups, have shown that p53 activity produces stress and context-dependent outputs ranging from selective gene regulation to metabolic inflammation and altered cell fate^48–50^. This perspective is relevant to vascular aging, where inflammatory programs may reflect organized tissue stress responses. Aging is accompanied by chronic, low-grade inflammation, commonly referred to as inflammaging, a concept linked to persistent sterile inflammation and cardiovascular, metabolic, and frailty-related phenotypes^51–53^. Chronic inflammation is also recognized as a hallmark of aging, and attenuation of persistent inflammatory signaling has been proposed as a strategy to improve age-associated tissue dysfunction^54^. Our findings refine this view in the context of vascular aging. The Aging_1 program, enriched for interferon-stimulated genes (ISGs), was defined from our single-cell analysis of ECs. When projected back onto aged tissues, it marked focal inflammatory sites organized around vascular structures, indicating spatially restricted vascular inflammation. Thus, even this broadly inducible IFN program is spatially organized around the aged vasculature. This finding supports a view of vascular inflammaging in which ECs not only are passive responders to systemic inflammation but are also spatial anchors around which immune and stromal inflammatory responses assemble occur. The coordinated enrichment of Irf7, Irf9, Stat1, and Stat2 suggests the activation of a type I IFN regulatory axis^55–57^. Sterile damage pathways such as cGAS–STING activation may link DNA damage, senescence, type I IFN induction, and aging-associated inflammation^58–61^.

Spatial transcriptomics clarified how IFN-responsive endothelial states are organized in aged tissues. Aging_1-high regions formed perivascular ISG clusters embedded in multicellular neighborhoods containing endothelial, stromal, myeloid, and mural cell populations. This organization is consistent with recent spatial work showing that vessel-associated niches are major hotspots of cardiac aging and inflammaging^46^. Our data extend this concept by identifying perivascular ISG clusters as local units of endothelial IFN activation in aged heart and adipose tissue. Importantly, these clusters were not simply sites of inflammatory accumulation. In aged adipose tissue, TNF-enriched immune aggregates were observed near ISG-positive vascular regions, which was consistent with the local inflammatory input. In parallel, spatial mapping of C3-related signals revealed that C3-expressing regions were positioned near Lrp1-positive areas, whereas C3ar1 exhibited a distinct distribution. Given the scavenger and endocytic functions of LRP1, this C3–Lrp1 spatial relationship may reflect local clearance or buffering within aging vascular niches^62,63^. Consistent with these findings, C3 depletion further increased IRF7 expression in aged hearts, suggesting that complement-related spatial signals may modulate perivascular IFN inflammation.

BST2 provides an example of how an endothelial IFN program can generate regulatory outputs in addition to inflammatory activation. In our functional assays, BST2 promoted a CD200R-associated macrophage state and reduced atherosclerotic lesion formation, suggesting that IFN-induced endothelial molecules can shape myeloid behavior in a protective direction. This concept is consistent with the broader principle that interferon-stimulated molecules can limit, as well as propagate, inflammatory responses^64^. Thus, endothelial BST2 may represent one downstream route through which chronic IFN activation is converted into macrophage-directed vascular protection.

Our study reframes vascular inflammaging as a local regulatory process, not a uniform inflammatory state. The conserved endothelial IFN response is one component of this process, with its consequences shaped by organ context, neighboring immune and stromal cells, complement-associated clearance signals, and endothelial effectors such as BST2. These findings argue against indiscriminate suppression of IFN signaling and instead support targeting specific endothelial–immune communication programs within diseased vascular niches. This perspective may help explain organ-selective vascular disease and guide more precise immune interventions in vascular aging and atherosclerosis.

### Limitations of the study

This study had several limitations. Multi-organ single-cell RNA-seq analysis was initiated with limited biological replicates per condition. Cell-level transcriptomic and cell–cell communication analyses were therefore used primarily as a discovery framework to nominate candidate endothelial stress programs and endothelial–immune interactions, which were prioritized through spatial transcriptomics, image-based validation, and functional perturbation. Further genetic and organ-specific perturbation studies will be needed to define the causal hierarchy of the C3- and BST2-associated pathways. Nevertheless, the validated findings support an IFN-centered link among endothelial stress responses, perivascular immune niches, and inflammatory remodeling in aged vasculature.

## RESOURCE AVAILABILITY

### Lead contact

Further information and requests for resources and reagents should be directed to and will be fulfilled by the lead contact, Tomoaki Tanaka.

### Materials availability

This study did not generate new unique reagents.

### Data and code availability

- The RNA-seq, single-cell RNA-seq, Visium HD spatial transcriptomic, and Xenium in situ gene expression datasets generated in this study have been deposited in the DNA Data Bank of Japan (DDBJ) Genomic Expression Archive (GEA). Accession numbers are E-GEAD-1267, E-GEAD-1268, E-GEAD-1269, E-GEAD-1270, and E-GEAD-1277.
- Raw sequencing reads linked to these GEA records have been deposited in the DDBJ Sequence Read Archive (DRA).
- The publicly available datasets used in this study are listed in the key resources table and in the Methods section.
- The custom code used to reproduce the transcriptomic analyses is provided in Supplementary_Code_Transcriptomic_Analyses.md.
- Any additional information required to reanalyze the data reported in this paper is available from the lead contact upon request.

## Acknowledgements

We thank Yoko Sasaki for mouse colony management and laboratory management, Mie Kobayashi for cryosectioning and laboratory management, Tsubasa Matsuzaka for paraffin sectioning suitable for Visium and Xenium analyses, and Dr. Masayuki Ota for assistance with block facing and section preparation for Visium analysis.

## Author contributions

M.Y. and T.T. designed the study. M.Y., A.N., Y.T. and M.C. performed the experiments, and M.Y., M.C. and M.S. analyzed the data. Y.G. prepared the RNA-seq libraries. M.F. contributed to single-cell analysis workflows and tool selection, and K.I. overviewed data analysis. J.I. provided the histological preparation assistance. M.Y. wrote the manuscript, and T.T. supervised the study.

## Declaration of generative AI and AI-assisted technologies

During the preparation of this work, the authors used ChatGPT to assist with editing the manuscript for English grammar, clarity, and readability, and to support the drafting and troubleshooting of analysis code by implementing established computational methods. All analyses, interpretations, and conclusions were reviewed and verified by the authors, who take full responsibility for the content of the published article.

## SUPPLEMENTAL INFORMATION

Document S1. Figures S1–S7 and supplemental figure legends

Data S1. Supplementary data file_regulonTargetsInfo, related to Figures 1–3

Data S2. Supplementary data file_scRNA analysis data, related to Figures 1–4

Data S3. Supplementary data file_Experimental raw data, related to Figures 5–7, S1

Supplementary_Code_Transcriptomic_Analyses.md. Custom code used to reproduce transcriptomic analyses, related to STAR Methods

## STAR★METHODS

### KEY RESOURCES TABLE

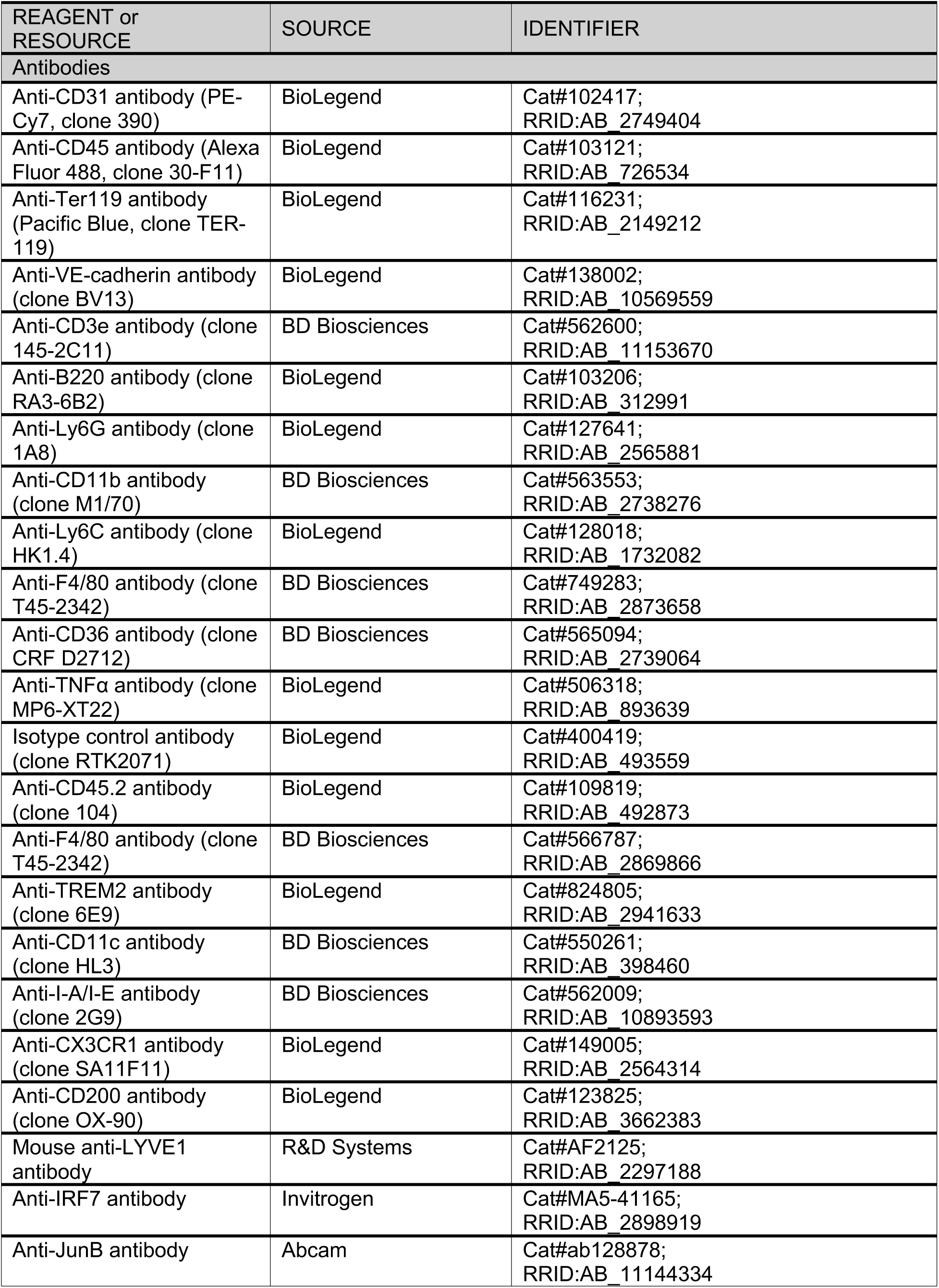
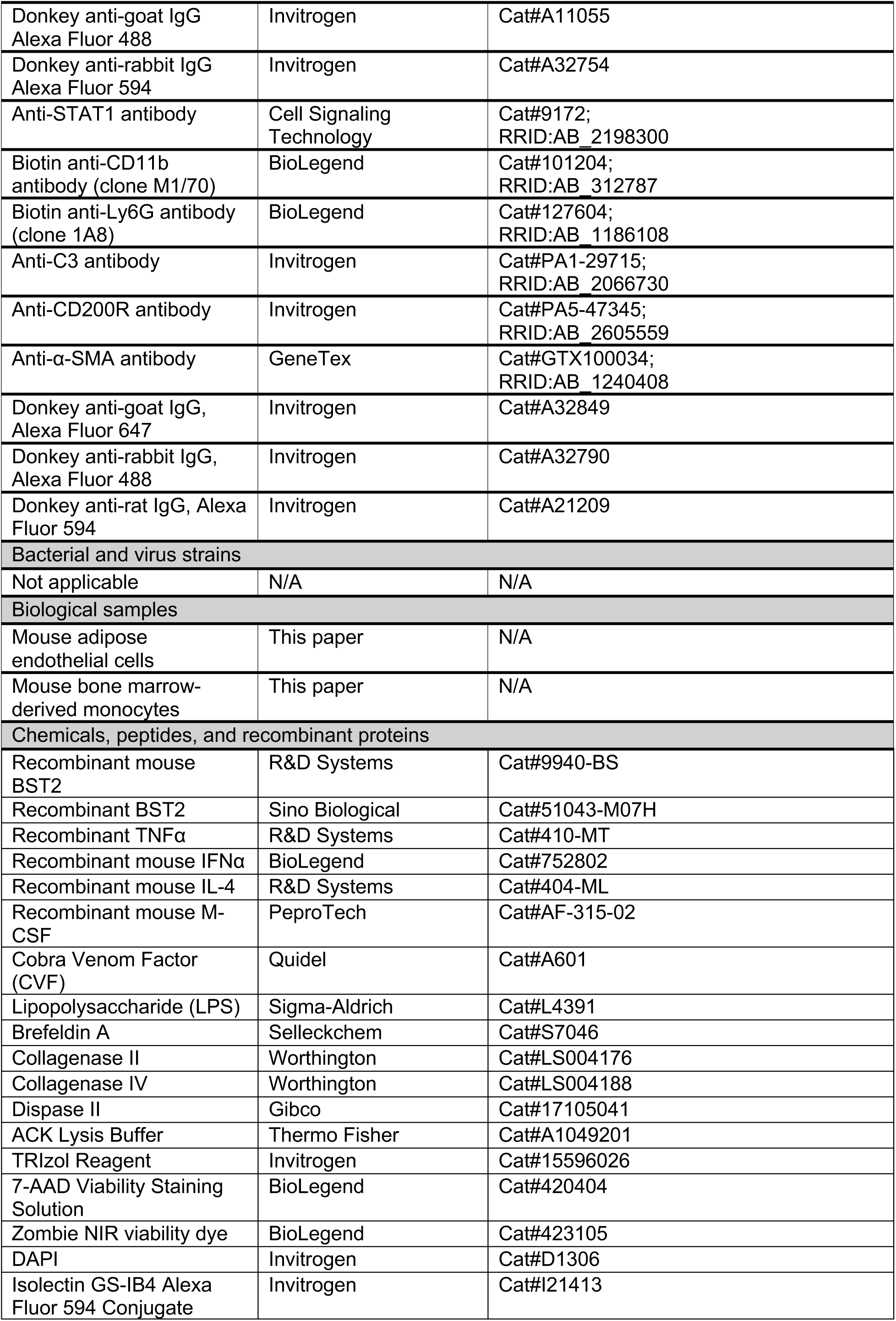
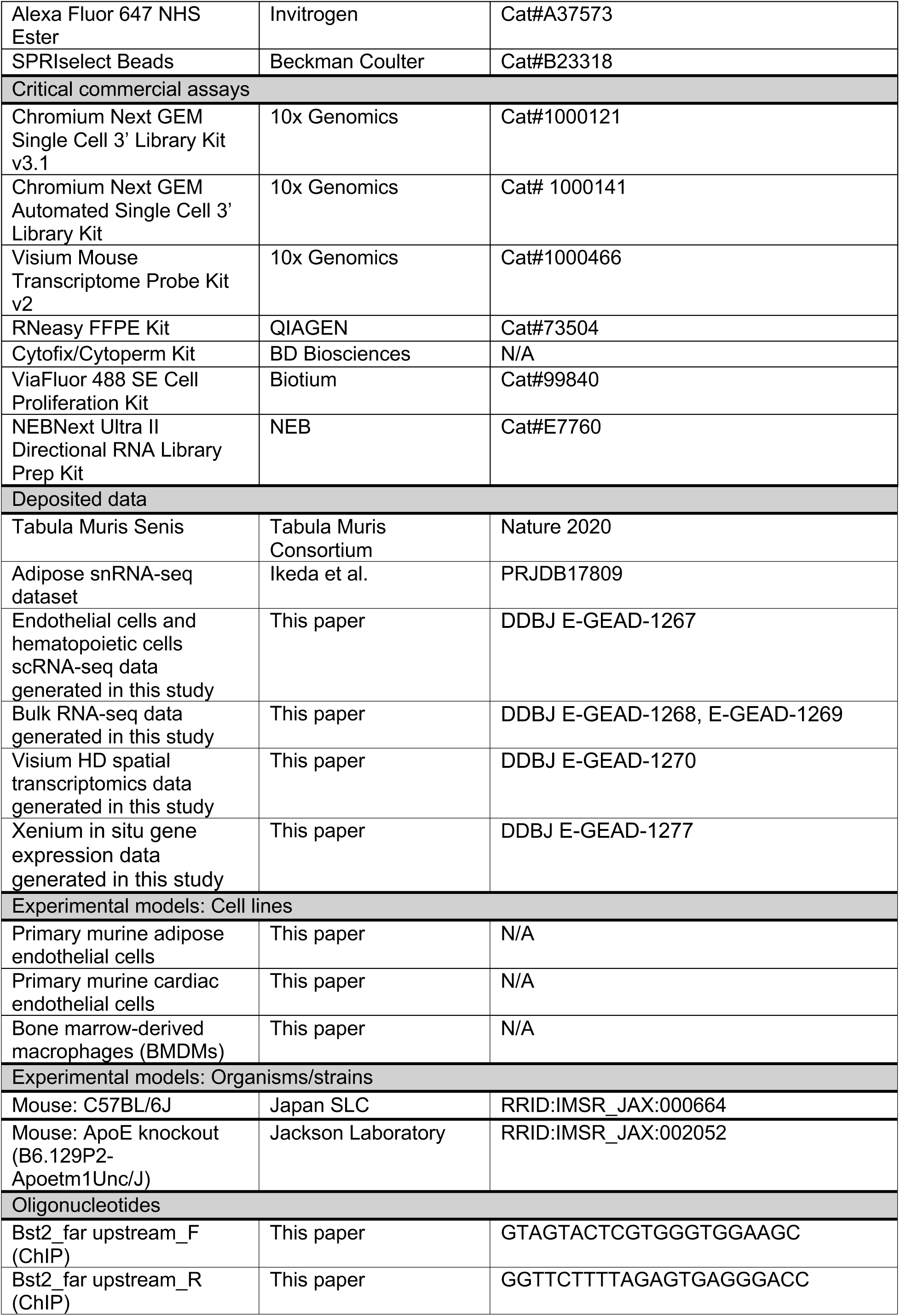
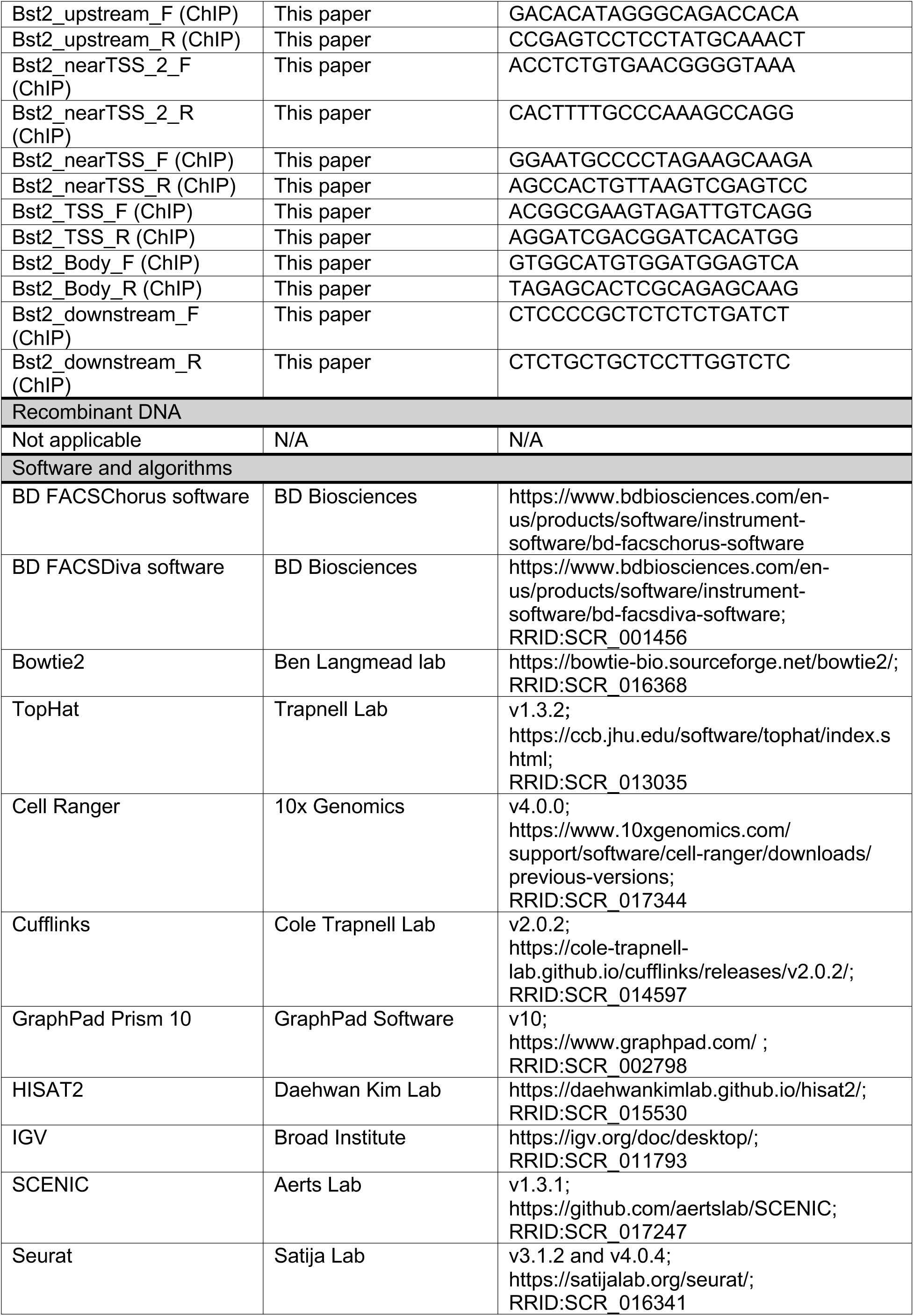
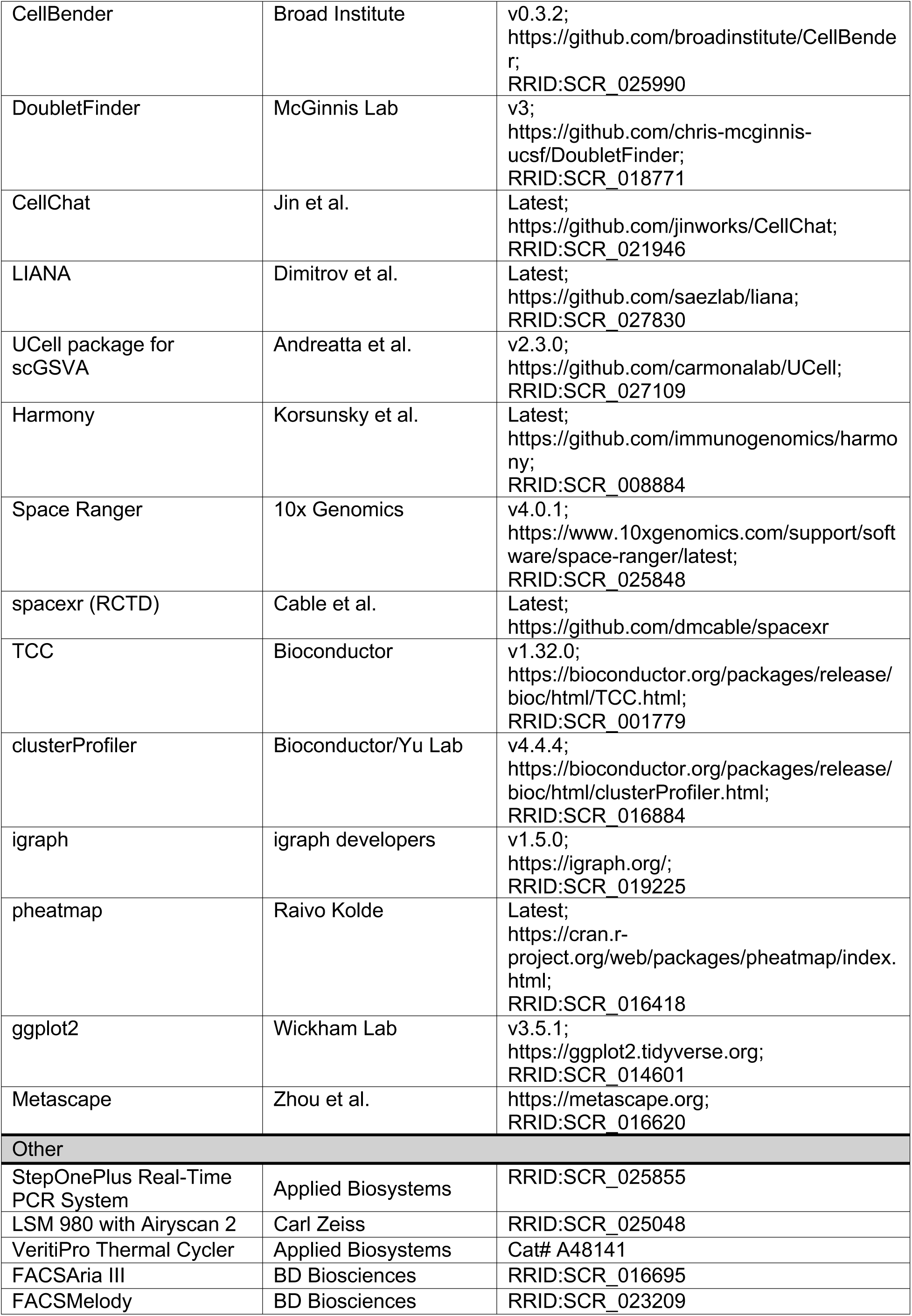
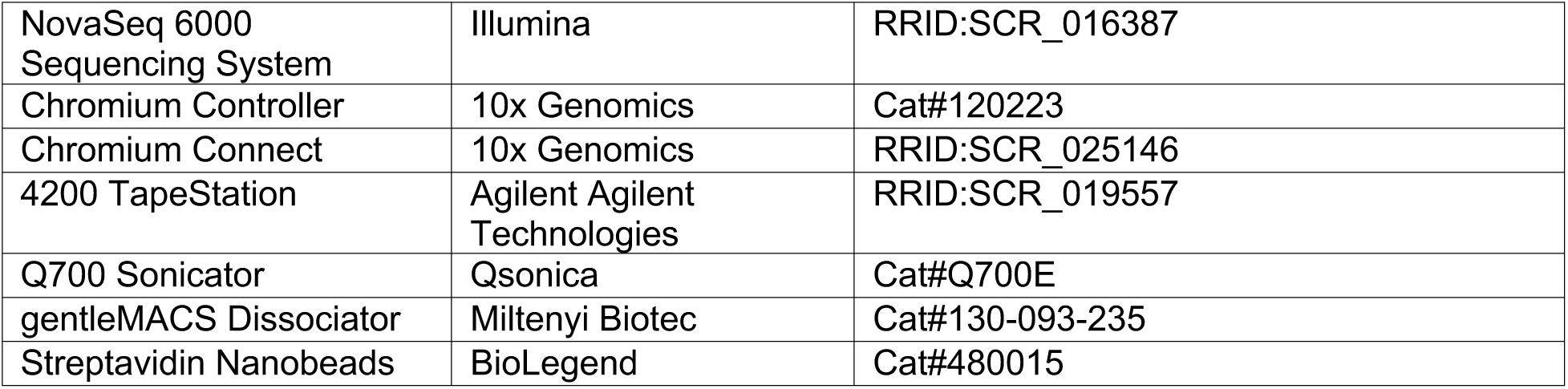

### EXPERIMENTAL MODEL AND STUDY PARTICIPANT DETAILS

#### Mice

All animal experiments were performed in accordance with institutional guidelines and were approved by the ethics committee for animals at Chiba University. The mice were maintained under specific pathogen-free conditions at 22 °C with a 12 h light/dark cycle, 40–70% humidity, and ad libitum access to food and water. Body weight was measured at the time of sample collection before tissue harvesting. Male C57BL/6J mice were purchased from Japan SLC, Inc. at 6 weeks of age and used for single-cell RNA-seq analyses. For aging comparisons, the mice were analyzed at 3 or 12 months of age. For diet-induced metabolic stress experiments, mice were fed either a normal diet or a high-fat diet containing 60% kcal from fat (Research Diets, D12492). For Visium HD spatial transcriptomic analyses, young and very aged mice, corresponding to 3 and 24 months of age, were analyzed. For atherosclerosis experiments, ApoE-deficient mice (B6.129P2-Apoe/J) were fed a high-cholesterol diet (Research Diets, D12079B) starting at 12–15 weeks of age. Recombinant BST2 was administered to ApoE-deficient mice as described in the Methods section. For complement depletion experiments, 15-month-old wild-type C57BL/6J mice were treated with cobra venom factor or PBS as described in the Methods section. Unless otherwise indicated, male mice were used for single-cell and spatial transcriptomic analyses. Both male and female mice were included in selected atherosclerosis and complement depletion experiments. Sex was not analyzed as a primary biological variable, which represents a limitation when generalizing the findings across sexes.

### METHOD DETAILS

#### Cell sorting for single-cell analysis

To isolate endothelial cells and hematopoietic cells, FACSMelodyCellsorter (BD Biosciences) was used. First, we performed intravital VEcadherin staining by injecting 25µg of anti-VE-cadherin antibody (Biolegend, clone BV13) conjugated to Alexa Fluor 647 NHS Ester (Thermo Fisher) retro-orbitally into mice under 3% isoflurane anesthesia, and after 10 min, the mice were euthanized. The samples were harvested after perfusion with 10 ml of PBS. Bilateral epididymal fat was defined as adipose tissue. Muscle tissue was taken bilaterally from the quadriceps, tibialis anteriors, extensor digitorum longus, soleus muscles and gastrocnemius muscles. For adipose tissues, and heart and liver samples, the minced tissue samples were incubated with a mixture of collagenase-II (2.5 mg ml⁻¹, worthington-biochem)/Dispase-II (2.5 ml⁻¹, gibco) mixture at 37°C for 30 min. For muscle tissues, a collagenase-IV (2.0 mg ml⁻¹, worthington-biochem)/ Dispase-II (0.7mg ml⁻¹, gibco) mixture was used, and the tissues were incubated at 37°C for 40 min. The cells were filtered through a 40 μm cell strainer and centrifuged at 500 x g for 5 min, followed by RBC lysis treatment with ACK buffer (Thermo Fisher) on ice for 2 min to generate a single-cell suspension. The single-cell suspension was first blocked with an antibody against CD16/32 (2.4G2) before antibody staining. The following antibodies were used for additional staining; anti-CD31(PE-Cy7, clone 390, Biolegend), anti-CD45(Alexa Fluor 488 Clone 30-F11, BioLegend) and anti-Ter119 (Pacific Blue, BioLegend). Dead target cells were defined as DAPI^+^ cells. After eliminating dead cells or Ter119^+^ cells, VE-cadherin and CD31 double positive cells were sorted as endothelial cells, and CD45^+^ cells were sorted as hematopoietic cells.

#### Single cell RNA sequence library preparation

The scRNA-seq libraries were prepared according to 10× Genomics specifications. Equal numbers of the sorted endothelial cells and hematopoietic cells were mixed for each library and loaded onto the 10x Genomics Chromium Controller or 10x Genomics Chromium Connect. Samples from the 1-year-old mice were loaded into the 10× Genomics Chromium Controller to generate GEM beads, and the Chromium Next GEM Single Cell 3’ Library & Gel Bead Kit v3.1 was used for reverse transcription, cDNA amplification, and library construction, according to the manufacturer’s instructions. Samples from 3-month-old mice were loaded into 10× Genomics Chromium Connect. 10× Genomics Chromium Next GEM Automated Single Cell 3’ Library and Gel Bead Kit were used to generate the libraries. The libraries were quantified and quality-assessed with an Agilent TapeStation. The resulting libraries were sequenced on a NovaSeq 6000 on a paired-end read flow cell with Novaseq 6000 reagent kit v1.5 (SP or S1). The sequencing data were subsequently processed with the Cell Ranger pipeline (7.0.0) to generate gene expression matrices for downstream analysis.

#### Single-cell RNA sequence data analysis

Background noise was removed from the data using CellBender v0.3.2 by inputting the raw_feature_bc_matrix.h5 files. All data analyses were conducted using Seurat version 5.0.3. Cells were selected on the basis of the following cutoffs: nFeature_RNA > 1000, nFeature_RNA < 5000, and percent.mt < 10. After selection, SCTransform was performed for normalization on each sample. Assuming a 7.5% doublet formation rate, the DoubletFinder (v3) algorithm was applied to computationally remove doublets before proceeding with downstream analyses. To combine multiple datasets, variable features were determined using the SelectIntegrationFeatures function. Genes were filtered to remove those commonly associated with mitochondrial genes, ribosomal RNA, pseudogenes, noncoding RNAs, and other genes of uncertain function. Specifically, genes matching the following patterns were excluded:"mt-", "MT-", "MMU", "SNO", "Gm", "Rn6", "RNA", "7SK", "SNORD", "SNORA", "SCARNA", "B3g", "Vmn", "Mir", "Rik", "Snora", "Snord", "LOC", "Rn4", "OTTMUSG", "Scarna", "Rnu", "Rmr", "Rpl", "Rps", "AW", "BC", "Malat1", "H2-", "B2m", "Actb", "AA", "AB", "AC", "AF", "AI", "AL", "AU", "AV", "AW", "AY" and "Hist". This filtering process was conducted to ensure that only biologically relevant genes were included in the downstream analysis. The molecular expression of the top 100 genes with the highest levels of expression in endothelial cells derived from four organs was identified using the FindAllMarkers function with the following parameters: only.pos = TRUE, min.pct = 0.25, and logfc.threshold = 0.25. To extract the top 100 differentially expressed genes in the aging or diet comparisons, we first applied PrepSCTFindMarkers. The FindMarkers function was subsequently used with the parameters logfc.threshold = 0.25 and recorrect_umi = FALSE for each organ. The top 100 genes were selected on the basis of the smallest adjusted p values (p_val_adj) for each organ. Gene ontology analysis was performed using the enrichGO function with the following parameters: OrgDb = org.Mm.eg.db, keyType = "ENTREZID", and ont = "ALL". The p-value adjustment method was set to “BH” (Benjamini–Hochberg), with a p-value cutoff of 0.05 and a q-value cutoff of 0.05. To calculate the signature score, we used a modified 2D scoring approach. Control genes were selected on the basis of expression levels and the percentage of cells in which each gene was detected. For each gene in the target list, a set of control genes with similar expression patterns was identified. The final signature score was calculated by subtracting the mean expression of control genes from the mean expression of the target gene set. The signature score was calculated using a custom function, calculate_signature_score, which first selects control genes using the get_controls function. For each cell, the mean expression of the target gene list was calculated and the mean expression of a set of control genes was subtracted. This approach ensures that the signature score is adjusted for cellular complexity, reducing the influence of overall transcriptional activity.

#### Single-cell RNA-seq preprocessing and cell annotation

After clustering, cell annotation was performed on the basis of canonical marker gene expression. Vascular endothelial cells were defined by *Pecam1* and *Cdh5* expression and distinguished from *Ptprc*-positive hematopoietic cells and *Col1a1/Pdgfra*-positive fibroblasts. Hematopoietic cells were annotated into macrophage/monocyte/DC populations (Macrophage lineage), T cell/NK cell/ILC populations (T-cell lineage), B cell/plasma cell populations (B-cell lineage), neutrophils, and mast cells. Representative markers used for annotation included *Cd68* and *Csf1r* for the Macrophage lineage; *Cd3e*, *Cd3d*, and *Nkg7* for the T-cell lineage; *Ms4a1* and *Cd19* for the B-cell lineage; *Ngp* and *S100a8* for neutrophils; and *Tpsb2* for mast cells. Clusters expressing markers of multiple lineages were designated as mixed cells, and proliferating cells were defined by *Mki67*, *Top2a*, and *Cdk1* expression. Mixed-lineage and proliferating cells were excluded from subsequent endothelial analyses. Vascular cells were subclustered to define endothelial and mural cell populations. Pericytes/vascular smooth muscle cells were annotated using *Pdgfrb*, *Cspg4*, *Tagln*, and *Acta2*. Lymphatic endothelial cells were annotated using *Prox1*, *Pdpn*, and *Lyve1*. In adipose tissue, skeletal muscle, and heart tissue, blood vascular endothelial cells were classified into arterial, venous, and capillary endothelial cells using the following established vascular markers: *Fbln5* and *Hey1* for arterial endothelial cells, *Vwf*, *Nr2f2*, and *Vcam1* for venous endothelial cells, and *Rgcc* and *Car4* for capillary endothelial cells. Liver endothelial cells were annotated separately. *Lyve1* and *Vwf* were used to distinguish liver sinusoidal endothelial cells from major-vessel endothelial cells, and *Vwf*-positive liver endothelial cells were further subdivided into central vein and portal vein endothelial cells using *Rspo3*, *Bmp4*, *Adgrg6*, and *Nrg1*. B-cell lineage cells were subdivided on the basis of canonical marker expression. Pro-B cells were defined by the expression of *Il7r*, *Rag1*, and *Myb*, whereas Ms4a1-positive B-cell clusters with high *Vpreb3* expression were designated pre-B cells. Mature and memory B cells were annotated using *Ighd*, *Fcer2a*, *Zbtb20*, *Zbtb32*, and *Bhlhe41*. Clusters that were *Cd19*-negative, *Ms4a1*-negative, and *Jchain*-positive were classified as plasma cells. For the T-cell lineage, *Cd3e* and *Cd3d* were used to define T cells. *Cd3*-negative cells expressing *Nkg7* and *Klrb1c* were classified as NK or ILC1 populations on the basis of *Gzma* and *Xcl1* expression, whereas ILC2 cells were identified by *Gata3*, *Il1rl1*, *Klrg1*, and *Rora*. Among Cd3-positive T cells, *Tmem176a* and *Tmem176b* expression defined Th17 cells, and the coexpression of *Trdc* and *Trdv4* marked γδTh17 cells. *Foxp3*-positive and *Ctla4*-positive clusters were annotated as regulatory T cells. *Cd4* and *Cd8a* expression distinguished *CD4*-positive and *CD8*-positive T cell subsets, with *Ccr7* and *Lef1* marking naïve T cells. For the Macrophage lineage, monocyte, macrophage, and dendritic cell subsets were annotated using canonical myeloid markers. Dendritic cell subsets (moDC: monocyte-derived, pDC: plasmacytoid, mRegDC: maturation-associated/Regulatory) were annotated using *Cd209a*, *Siglech*, *Ccr9, Ccr7*, and *Il4i1*. *Plac8*-positive cells were designated monocytes and were further separated by *Ly6c2*, *Chil3*, *Ace*, and *Ccr2* expression. Macrophage subsets were annotated using *Cd163*, *Retnla*, *Ear2*, *Ccr2*, *Cx3cr1*, and *Trem2*. Kupffer cells were identified as a liver-specific macrophage population coexpressing *Cd5l* and *Clec4f*. The relative abundance of these immune subsets were compared across organs and conditions to support source-cell assignment in the CCI analysis.

#### SCENIC analysis for transcription factor network characterization

Transcription factor–target regulatory networks were inferred using single-cell regulatory network inference and clustering (SCENIC) implemented in R (v1.3.1). Two motif databases were used: “mm10 refseq-r80 10kb_up_and_down_tss. mc9nr.feather” and “mm10 refseq-r80 500bp_up_and_100bp_down_tss.mc9nr. feather.” Regulons were identified under the *extended* mode, which incorporates both high-confidence and low-confidence motif annotations (highConfAnnot = TRUE/FALSE). For each organ dataset, regulon activity per cell was quantified using the *regulonAUC* function, and the regulon specificity score (RSS) was computed with *calcRSS*. Regulons with an RSS > 0.01 and Z > 1.0 were retained for subsequent analyses. Hierarchical clustering of regulons was based on *Euclidean* distance and *ward.D* linkage, and correlation analyses were performed using the *Spearman* method. Organ-specific and condition-specific comparisons were conducted across multiple analytical tiers. For aging analysis, SCENIC was performed independently on endothelial subsets from young and aged mice fed a normal diet. Regulons were ranked by their RSS difference, and the top transcription factors enriched in aged samples were selected for each organ. These organ-level results were merged to identify shared and tissue-restricted aging-responsive regulons. The coexpression strength of the coexpression between each regulon and its target genes was obtained from *regulonTargetsInfo* (CoexWeight parameter) and used to define aging-related regulatory clusters. For dietary stress analysis, the same procedure was applied to endothelial cells from Normal diet or a HFD. Differentially enriched regulons were ranked, and cross-organ comparisons were used to delineate diet-responsive regulatory modules. For integrative assessment across all conditions (four organs × four states: Young/Normal, Aged/Normal, Young/HFD, Aged/HFD), regulonAUC and RSS were calculated for each dataset, enabling the comparison of regulon activity across stress types and tissues. Correlation matrices were constructed using *cor(method = "spearman")* and *hclust(method = "ward.D2")* to identify coregulated modules. For gene–regulon module analysis, the union of the top stress-responsive genes from each organ was combined with the union of the top stress-responsive regulons identified by SCENIC. Coexpression weights between each regulon and its target genes were extracted from regulonTargetsInfo using the CoexWeight parameter and assembled into a gene–regulon matrix. Hierarchical clustering of this matrix was used to define stress-responsive gene–regulon modules. To evaluate whether the identified regulons were associated with the corresponding gene modules at the single-cell level, module signature scores were calculated for each gene module using expression-matched control genes. SCENIC-derived regulonAUC values were subsequently compared with the corresponding module signature scores in individual endothelial cells. Pearson correlation coefficients were calculated for each regulon–module pair and summarized to identify regulons associated with shared or organ-biased stress-responsive modules.

#### Correlation analysis between transcription factor regulon activity and pathway enrichment

To investigate how stress-responsive transcription factor (TF) regulons are functionally linked to cellular signaling states, we assessed the correlation between regulon activity and pathway enrichment scores at the single-cell level. For each organ (adipose tissue, skeletal muscle, liver, and heart), endothelial cells were subsetted from the Seurat object and analyzed separately under aging and diet conditions. For each comparison, TF regulons showing stress-associated changes in SCENIC regulon activity were selected for subsequent pathway correlation analysis. Pathway activity was quantified using single-cell gene set variation analysis (scGSVA) implemented with the UCell package (v2.3.0) and Reactome pathway annotations. The buildAnnot() function was used to construct the Reactome reference matrix with species = “mouse”, keytype = “SYMBOL”, and anntype = "Reactome". Enrichment scores were computed using scgsva() with method = "UCell" and maxRank = 5000. The resulting pathway scores were integrated into the Seurat object metadata for downstream analysis. For each selected TF regulon, regulon activity values were extracted from the SCENIC regulonAUC_extended assay, and Spearman correlation coefficients were calculated between regulon activity and Reactome pathway enrichment scores across single endothelial cells within each organ and stress comparison. Only positive correlations were retained for downstream ranking. The squared Spearman correlation coefficient was calculated as R² and used to evaluate the strength of association. Pathways with R² > 0.1 were considered correlated with the corresponding regulon. The top 20 positively correlated pathways for each regulon were summarized and visualized using ggplot2 (v3.5.1), ranked by R². To examine the global relationships between TF regulons and pathways, correlation networks were constructed by linking each TF regulon to its top correlated pathways across organs. Nodes representing TF regulons and pathways were connected by weighted edges according to their Spearman correlation coefficients and visualized using igraph (v1.5.0). Nodes corresponding to pathways or TF regulons with correlation coefficients greater than 0.3 were selectively labeled, and their node borders were emphasized to highlight highly connected relationships. Similarly, edges with correlation coefficients greater than 0.3 were drawn with increased thickness compared with the baseline edges, emphasizing stronger associations within the transcription factor–pathway network.

#### Cell–cell communication inference

Intercellular signaling was inferred with CellChat (mouse database) on SCTransform-normalized counts for four organs (adipose, heart, skeletal muscle, and liver) under four conditions (young vs. aged; normal diet vs. high-fat diet), yielding 16 independent analyses saved as RDS objects for downstream comparison. For each organ–condition dataset, we constructed a CellChat object using curated broad cell classes as grouping variables (capillary, arterial, and venous endothelium; T/NK/ILC; B/plasma; macrophage/monocyte/DC). The standard pipeline was executed in sequence: normalizeData, subsetData (to signaling genes in CellChatDB.mouse), identifyOverExpressedGenes and identifyOverExpressedInteractions with package defaults, optional projection onto the mouse PPI network (projectData), computeCommunProb (mass-action formulation of ligand–receptor usage with built-in permutation-based significance), filterCommunication (min.cells = 1), computeCommunProbPathway, and aggregateNet. Global summaries were visualized with netVisual_circle using interaction weights/strengths; these are the values reported in the main figures. Condition contrasts were performed pairwise within each organ using rankNet(mode = "comparison", stacked = TRUE, do.stat = TRUE) to extract pathways with significant changes in information flow between conditions (aging and diet comparisons). For endothelial-centric summaries, we focused on capillary endothelial cells and quantified incoming (indegree) and outgoing (outdegree) centrality per pathway from the pathway-level networks (@netP). To accommodate zero inflation across conditions, we reported condition-to-condition changes as ratios, log2-fold changes, and absolute differences for indegree, outdegree, and their sums (incoming + outgoing). Capillary EC–focused scatter plots were generated with netAnalysis_signalingChanges_scatter (idents.use = "Capillary Endothelial cell"). To map the differentially used ligand–receptor pairs, identifyOverExpressedGenes was rerun in the comparison objects with group.dataset = “datasets”, pos.dataset set to the stressed condition (e.g., "Normal_Aged" or "HFD_Aged"), and thresholds thresh.pc = 0.1, thresh.fc = 0.1, thresh.p = 1 to collect broad candidate sets, which were then intersected with inferred communication edges via netMappingDEG. Pathway subsets used in ratio/log2FC/difference summaries were curated by excluding ubiquitous junctional modules (e.g., PECAM1, ESAM, PTPRM, CDH5, MPZ, and KIT) and retaining stress-responsive families (e.g., MIF, CXCL, JAM, SEMA6/7, and LAMININ). All analyses were performed in R with fixed random seeds; the CellChat permutation framework was used for edge-level significance and pathway-level differential testing as implemented in the package.

#### Integration of CellChat-inferred signaling with transcriptional regulons and pathway activity

To link endothelial communication dynamics to transcriptional control and cellular functions, we integrated CellChat outputs with SCENIC regulon activity and per-cell pathway scores from UCell. Regulon activity was quantified using the SCENIC regulonAUC in extended mode on organwise single-cell objects; organ- and stress-specific TFs were taken from the discovery stage of the study and used for downstream association tests. Per-cell pathway activity was computed with UCell (Reactome mouse gene sets, SYMBOL keys) using maxRank = 5000; the resulting scores were stored in Seurat meta.data for joint analyses. For each TF, we computed Spearman correlations between its regulonAUC and all Reactome pathway scores across cells in the relevant organ–condition dataset, retained the top 20 pathways per TF, and summarized them in network graphs (igraph, Fruchterman–Reingold layout). Nodes represented TFs or Reactome pathways; and the edges were weighted by correlation coefficients. To aid interpretation, only nodes with correlation > 0.3 (together with all TF nodes) were labeled, and edges with correlation > 0.3 were rendered thicker (edge width = 3). In complementary gene-level analyses linking CCI edges to TF control, we intersected endothelial ligand/receptor genes present in CellChat pathways with SCENIC regulonTargetsInfo (coexpression weights and motif evidence, including highConfAnnot, nMotifs, bestMotif, NES, and coexpression statistics) and then filtered these genes by membership in organ-specific Reactome pathways compiled from the per-organ Reactome annotations. Sankey diagrams (ggalluvial) summarized Tissue → Gene → TF relationships after this triage. To quantify TF–gene coupling at single-cell resolution, we fit per-cell linear models (gene expression ∼ regulonAUC) in each organ and reported the R²; gene–TF pairs with R² > 0.08 were retained and ranked across all organs. For cross-organ summaries, the bar plots are colored by organ and restricted to pairs passing the R² cutoff. Throughout the study, the same cutoffs and display rules were applied uniformly across organs and comparisons to maintain comparability: UCell maxRank = 5000; top-20 pathways per TF by Spearman correlation; node labeling for correlation > 0.3; thick edges for correlation > 0.3; and R² > 0.08 for TF–gene regressions. The figures pooling capillary EC analyses across organs used axes ranges and aesthetics identical to those used in the CellChat scatter, ensuring that shifts in incoming/outgoing influence and their TF–pathway couplings could be compared directly.

#### Visium HD spatial transcriptomics library preparation

Mouse hearts and visceral adipose tissues were harvested and fixed in 10% neutral-buffered formalin (NBF). The samples were dehydrated through graded ethanol solutions, cleared in xylene, and embedded in paraffin wax (Tissue-Tek Paraffin Wax II 60, Sakura Finetek). FFPE blocks were cut into 10 µm thick sections. For RNA quality control, RNA was extracted from deparaffinized sections using an RNeasy FFPE Kit (QIAGEN, 73504), and RNA quality was assessed using an Agilent TapeStation. Samples with DV200 values greater than 30% were used for library preparation. Visium HD spatial transcriptomics libraries were prepared according to the manufacturer’s protocol using the Visium Mouse Transcriptome Probe Kit v2. Libraries were sequenced on an Illumina NovaSeq 6000 system with paired-end reads using an S1 flow cell.

#### Reference construction for spatial cell-type annotation

Tissue-specific reference datasets were generated for spatial cell-type annotation. Public mouse aging atlas datasets were obtained from CZ CELLxGENE Discover as h5ad files, including heart, adipose tissue, and gonadal fat pad datasets from the Tabula Muris Senis atlas, were obtained from CZ CELLxGENE Discover as h5ad files. The cell category columns were converted to string format before downstream processing. For the heart analysis, the reference dataset was generated by merging the Tabula Muris Senis heart dataset with the single-cell RNA-seq dataset generated in the present study, followed by normalization and integration. The resulting heart reference object was saved as reference_rctd_heart.rds and used for RCTD-based annotation with spacexr. For adipose tissue, the reference dataset was generated by merging the Tabula Muris Senis adipose tissue and gonadal fat pad datasets with our previously analyzed adipose single-nucleus RNA-seq dataset deposited under BioProject accession number PRJDB17809^33^. The resulting adipose reference object was saved as adipose_merged_obj_reference.rds and used for anchor-based label transfer to the Visium HD segmented adipose dataset using Seurat FindTransferAnchors and TransferData.

#### Visium HD data processing and cell-type annotation

The raw sequencing data were processed using Space Ranger v4.0.1. Heart samples were analyzed using 8 µm binned output matrices, whereas visceral adipose tissue samples were analyzed using cell segmentation-based output matrices. Downstream analyses were performed using Seurat v5.0.3. For each tissue, four samples were loaded as on-disk matrices using BPCells and converted into Seurat objects. Low-depth spatial units were removed using an nCount_RNA cutoff of 25. The datasets were normalized, variable features were identified, and sketch-based dimensional reduction was performed using 50,000 cells selected by leverage score. Principal component analysis was performed on the sketch assay, followed by Harmony-based integration using dimensions 1–30, UMAP visualization, and clustering. The integrated sketch analysis was then projected back to the full dataset.

Gene identifiers were converted from Ensembl IDs to MGI gene symbols using biomaRt, while unmapped Ensembl IDs were retained. For heart samples, RCTD was performed using spacexr on spatial units with nCount_RNA ≥30. Sample-specific spatial coordinates were obtained from tissue position files and matched to the corresponding 8 µm barcodes. RCTD was run in doublet mode using the heart reference dataset. RCTD labels classified as “reject” were assigned as “Unknown.” Final heart cell-type labels were generated by integrating marker-based cluster annotation, RCTD prediction, and manual curation of anatomical regions, including cardiac valve regions.

For adipose tissue, cell-type labels were transferred from the adipose reference dataset using Seurat anchor-based label transfer. Cells with a maximum prediction score less than 0.4 were assigned as “Unknown.” Ambiguous endothelial, myeloid, and unknown populations were further resolved by Harmony SNN-based reclustering and marker-based annotation. The final labels were merged into a simplified cell-type annotation. For adipose segmented cells, segmentation areas and centroid coordinates were extracted from Space Ranger cell segmentation GeoJSON files and added to the Seurat metadata for spatial visualization and distance-based analyses. Images of H&E-stained cells generated during Visium HD processing were visualized using Loupe Browser v9.0.0. Spatial gene expression overlays for *Tnf*, *Cxcl13*, *Irf7*, *C3*, *Apoe*, *Lrp1*, and *C3ar1* were generated using the same normalization and color-scale settings across compared samples.

#### Spatial cell–cell interaction analysis using LIANA

Spatial cell–cell interaction analysis was performed using LIANA. An interferon-related Aging_1 module score was calculated using Seurat AddModuleScore with a predefined interferon-stimulated gene signature. In heart samples, analyses were performed in left ventricular regions of interest, and cells with an Aging_1 score ≥0.3 were defined as core cells. In adipose tissue, cells with an Aging_1 score ≥0.05 were defined as core cells. Noncore cells located within 100 µm of the core cells were defined as neighboring cells, and all remaining cells were defined as distal cells. Spatial interaction groups were assigned as Core, Neighbor_, or Distal_ according to their spatial relationship and final cell-type annotation.

For LIANA input, normalized expression matrices and metadata containing spatial interaction groups were exported from Seurat objects. Heart analyses used the Spatial.008um data layer, whereas adipose analyses used the SCT data layer. Exported matrices were loaded into Python as AnnData objects, and LIANA rank aggregation was performed using the mouse consensus ligand–receptor resource with the CCI_group as the grouping variable. Ligand–receptor interactions targeting core cells were extracted from neighbor and distal source groups. For each source cell type and ligand–receptor pair, mean lrscore values were calculated separately for neighbor-to-core and distal-to-core interactions. Neighbor-enriched interactions were retained when the mean neighbor-to-Core lrscore exceeded the corresponding distal-to-Core lrscore. Results were summarized across two Young and two Aged samples, and ligand–receptor interactions reproducibly detected in both biological replicates of at least one age group were retained for downstream visualization.

#### Xenium in situ gene expression profiling

Mouse hearts, quadriceps femoris muscles, and visceral adipose tissues were processed for FFPE embedding as described above. Section quality was assessed by DAPI staining and Gapdh in situ hybridization, and additional 5 µm FFPE sections were mounted onto Xenium slides (PN-1000465; 10x Genomics). Deparaffinization and decrosslinking were performed according to the Xenium In Situ for FFPE Tissue Preparation Guide (CG000578 Rev F; 10x Genomics).

In situ gene expression profiling was performed using the Xenium platform (10x Genomics). The Xenium Mouse Tissue Atlassing Panel v1, containing 379 genes, was supplemented with a 100 gene custom add-on panel designed using the Xenium Panel Designer (design ID: BE76YW; BE76YW_mMulti_100g). The custom panel included genes from the Aging_1 to Aging_4 modules, together with annotation markers for inflammatory cells and endothelial cells identified from our single-cell RNA-seq analysis.

Probe hybridization, ligation, amplification, cell segmentation staining, autofluorescence quenching, imaging, and decoding were performed according to 10x Genomics Xenium protocols. Slides were processed on the Xenium Analyzer using instrument software version 4.0.1.4 and analysis software version 4.0.1.0. After the Xenium run, H&E staining was performed on the same slides. Xenium output files were evaluated using Xenium Explorer version 4.1.1, and H&E images were aligned to Xenium data. Quality control was performed by reviewing the Xenium Analyzer summary HTML files and confirming that no run-level errors were reported. Aging_1 scores were calculated from the Aging_1 gene set, and cell segmentation outputs were used to define cellular compartments and map target gene expression. Xenium cell boundaries were generated using the default multimodal segmentation workflow, which integrates DAPI-based nuclear segmentation, interior RNA staining with a universal 18S rRNA label, and antibody-based boundary staining for ATP1A1, E-cadherin, and CD45.

#### Analysis of Junb regulon activity in adipose endothelial cells

To investigate stress-responsive transcriptional programs in adipose endothelial cells, we focused on *Junb* activity inferred from SCENIC regulon analysis. Endothelial cells from adipose tissue under normal diet conditions were reclustered, and their *Junb* regulon activity scores (RegulonAUC, extended mode) were determined. Cells were dichotomized into “Junb high” and “Junb low” groups using a cutoff value of 0.15, corresponding to the median range of the regulon activity distribution. Within these subsets, the samples were further divided by age (young and aged), generating four subgroups for comparison.To identify genes associated with *Junb*-dependent transcriptional activation during aging, differential expression analysis was performed between the “Junb high” and “Junb low” subsets within the aged group using the FindMarkers function in Seurat (log₂FC threshold = 0.1). The top 100 differentially expressed genes were selected on the basis of the highest average log₂ fold change.These candidate genes were subsequently subjected to functional enrichment analysis using Metascape (www.metascape.org), and upstream transcriptional regulators were ranked using the TRRUST database to identify key factors potentially mediating *Junb*-associated transcriptional programs in aged adipose endothelial cells.

#### Immunofluorescence staining of frozen sections

Intravital VE-cadherin staining was performed by retro-orbital injection of 25 µg of anti-VE-cadherin antibody (BioLegend, clone BV13) preconjugated to Alexa Fluor 647 dye (Invitrogen, A37573) into mice under 3% isoflurane anesthesia. The mice were euthanized 10 min after injection, and the organs were harvested and fixed overnight in 4% paraformaldehyde (PFA) at 4 °C. For immunostaining, the fixed samples were cryoprotected in 30% sucrose overnight following PFA fixation, embedded in OCT compound, and sectioned at a thickness of 5 µm. For C3 staining, fresh tissues were embedded in OCT compound without prior fixation, frozen, sectioned, and fixed with cold methanol at −20 °C for 10 min. The tissue sections were permeabilized either with PBS containing 0.1% Triton X-100 for 10 min at room temperature or with cold methanol at −20 °C for 10 min. Blocking was performed with PBS containing 1% bovine serum albumin (BSA; Thermo Fisher, 37525) and 5% normal donkey serum for 1 h at room temperature. Immunofluorescence staining was carried out using the following primary antibodies: anti-LYVE1 (AF2125, 1:100), anti-IRF7 (MA5-41165, 1:100), anti-JunB (ab128878, 1:100), anti-C3 (Invitrogen, PA1-29715, 1:500), anti-CD200R (Invitrogen, PA5-47345, 1:50), anti-F4/80 antibody (BD Biosciences, clone T45-2342, 1:50), and anti-α-SMA (GeneTex, GTX100034, 1:1000). The secondary antibodies used were anti-goat IgG conjugated with Alexa Fluor 647, anti-rabbit IgG conjugated with Alexa Fluor 488, and anti-rat IgG conjugated with Alexa Fluor 594 (Invitrogen). AF594-conjugated Isolectin IB4 (Invitrogen, I21413) was also used to stain vascular endothelial cells. Nuclear counterstaining and mounting were performed using Mounting Medium with DAPI (VECTASHIELD, Vector Laboratories). Fluorescence images were acquired using a Zeiss LSM 980 confocal microscope. The IRF7-positive area was quantified using Fiji/ImageJ. IRF7-positive pixels were quantified using a fixed threshold applied uniformly to all the images. For each mouse, three fields were analyzed, yielding a total of 12 fields from four mice.

#### Flow cytometry analysis of macrophage populations in tissues

The mice were fed either a normal diet or HFD from 6 weeks of age and analyzed after 15 months of feeding. Tissue dissociation was performed using a mixture of collagenase and dispase optimized for each organ, as indicated in the preparation for single-cell analysis. Samples were digested using a gentleMACS Dissociator (Miltenyi Biotec) with the following protocols: 37C Multi_G for heart, 37C mLIDK1 for liver, 37C mrSMDK1 for skeletal muscle, and 37C mrATDK1 for adipose tissue. The digested suspensions were resuspended in 37% Percoll density gradient medium (Cytiva), and the cells were collected from the interface between the 37% and 70% Percoll layers. Flow cytometric analysis was performed on a FACSymphony A3 (BD Biosciences). Cells were stained with the antibodies listed in the antibody table, and live cells were gated as 7-AAD⁻. The macrophages were defined as CD31⁻CD45⁺B220⁻CD3e⁻Ly6G⁻CD11b⁺F4/80⁺, and the subpopulations were evaluated on the basis of Ly6C expression levels.

#### Flow cytometry analysis of TNFα levels in adipose macrophages

Intracellular cytokine production was measured following Golgi retention using Brefeldin A (Selleckchem, S7046), as previously described^65^. A 500 µg ml⁻¹, PBS solution of brefeldin A was intraperitoneally administered to the mice, and visceral adipose tissue was collected 6 h later for analysis. Comparisons were made between 8–10-week-old and 40-week-old mice. Adipose tissue was digested with collagenase II/Dispase II solution using a gentleMACS Dissociator (Miltenyi Biotec) according to the 37C mrATDK1 protocol, for single-cell preparation. Cells were isolated from the interface between the 37% and 70% Percoll layers (Cytiva). For Ly6G⁺ cell depletion, biotin-conjugated anti-Ly6G antibody (BioLegend, clone 1A8; 5 µL per sample) was added to the cells, which were incubated on ice for 15 min, followed by washing and the addition of streptavidin (SAV) beads (BioLegend; 5 µL). After a further 15-min incubation on ice, the samples were diluted to 2 ml with buffer and subjected to magnetic depletion using an EasySep instrument (STEMCELL Technologies). The remaining cells were stained for surface markers on ice for 30 min, fixed and permeabilized with a Cytofix/Cytoperm kit (BD Biosciences) for 40 min on ice, and then incubated overnight with an anti-TNFα antibody (BioLegend, clone MP6-XT22). Flow cytometric analysis was performed using a FACSymphony A3 (BD Biosciences). Zombie NIR viability dye (BioLegend) was included during surface staining, and dead cells were excluded together with CD3e⁺/B220⁺ populations using the APC-Cy7 gate.

#### Coculture of adipose endothelial cells with macrophages

Murine adipose endothelial cells were isolated using an anti-CD31 antibody (BioLegend, clone MEC13.3; 4.8 µL) preconjugated to anti-rat IgG Dynabeads (Invitrogen, 11036). The conjugated beads were incubated with digested adipose tissue for 30 min to capture CD31⁺ endothelial cells. Cells were cultured in endothelial growth medium consisting of DMEM/F-12 (400 ml), FBS (100 ml), HEPES (10 ml), heparin stock (SIGMA, 10 mg ml⁻¹; 5 ml), NEAA (5 ml), penicillin/streptomycin (5 ml), and Endothelial Cell Growth Supplement (Corning, 25 mg ml⁻¹; 1 ml). At passage 4, 3.6 × 10⁴ endothelial cells were plated per well in 24-well plates. Macrophages were isolated from the adipose tissue of 8-month-old mice. Tissue digestion and Percoll separation were performed as described for TNFα analysis. For Ly6G⁺ cell depletion, biotin-conjugated anti-Ly6G antibody (BioLegend, clone 1A8; 5 µL) and streptavidin (SAV) beads (BioLegend; 5 µL) were added sequentially. The remaining cells were then incubated with an anti-CD11b antibody (BioLegend, clone M1/70; 5 µL) and SAV beads (BioLegend; 5 µL) to enrich macrophages. After surface staining with the antibodies listed in the antibody table, the macrophages were sorted on a FACS Melody Cell Sorter (BD Biosciences) as 7-AAD⁻ CD45⁺ Ly6G⁻ CD11b⁺ F4/80⁺ cells and further separated into Trem2⁻ CD36⁻ and Trem2⁺ CD36⁺ populations. For co-culture, sorted macrophages (1.0 × 10⁴ cells) were seeded in the upper chamber of a Cell Culture Insert (BD, 353097; 24-well, 8.0 µm pore size) and co-cultured with endothelial cells in the lower chamber. M-CSF (20 ng ml⁻¹ M-CSF (PeproTech, AF-315-02) was added during the co-culture period. To examine the direct effects of TNFα, recombinant TNFα (R&D Systems, 410-MT) was added at 100ng mL⁻¹ to endothelial cells in parallel. Endothelial cells were harvested after 18 h of co-culture or 6 h of TNFα treatment for RNA-sequencing analysis.

#### ChIP analysis of STAT1 binding to the *Bst2* locus

Murine cardiac endothelial cells were isolated using an anti-CD31 antibody, following the same procedure as that for adipose endothelial cells, and cultured in endothelial growth medium. At passage 3, cells obtained from one heart were expanded in six 10-cm dishes until they reached confluence and then divided into control and stress groups. For the stress condition, recombinant mouse IFNα (BioLegend, 752802; 100 ng ml⁻¹) was added for 6 h, followed by the addition of LPS (Sigma-Aldrich, L4391; 100 ng ml⁻¹) for an additional 4 h. Cells were crosslinked with 1% formaldehyde at 37 °C for 10 min, and crosslinking was quenched with glycine at a final concentration of 0.125 M. The cells were subsequently lysed in swelling buffer (25 mM HEPES, pH 7.9; 1.5 mM MgCl₂; 10 mM KCl; and 0.1% NP-40) containing a protease inhibitor cocktail (Nacalai Tesque). Nuclei were collected by centrifugation, resuspended in lysis buffer (50 mM Tris-HCl, pH 8.0; 1% SDS; and 10 mM EDTA), and sonicated using a Q700 sonicator (Qsonica, Newtown, CT, USA) at 60% amplitude with 15 s on/45 s off pulses, until a total energy input of 6,000 J was reached. Ten percent of the chromatin was saved as input, and the remaining material was incubated with antibody-bound beads. Anti-STAT1 antibody (Cell Signaling Technology, #9172) was preconjugated to Dynabeads Protein G (Thermo Fisher Scientific) and added to the diluted chromatin, followed by incubation at 4 °C overnight with rotation. After being washed, the bound complexes were eluted twice in ChIP elution buffer (50 mM Tris-HCl, pH 7.5; 1 mM EDTA; and 1% SDS) at 65 °C for 5 min and then rotated at room temperature for 15 min. NaCl and RNase A were added to final concentrations of 160 mM and 20 µg ml⁻¹, respectively, and the samples were incubated overnight at 65 °C to reverse the crosslinks. Proteinase K and EDTA were then added to final concentrations of 200 µg ml⁻¹ and 5 mM, respectively, followed by incubation at 45 °C for 2 h. DNA was subsequently purified using a PCR purification kit (Viogene BioTek, New Taipei City, Taiwan). 1 µL of immunoprecipitated or input DNA was used as template for qPCR with the primer sets listed in the Primer List, and enrichment was quantified as a percentage of the input.

#### Bone marrow–derived macrophage (BMDM) differentiation with BST2 stimulation

Bone marrow cells were collected from both the tibiae and the femora of 6–8-week-old mice. Littermates were used for each assay. The procedure was adapted from a published protocol^66^ with minor modifications. Bone marrow was flushed from the bones using PBS and a 26G needle with syringe, followed by centrifugation at 350 × g for 4 min. The cell pellet was resuspended in 1 ml of ACK buffer (Gibco, A1049201) for red blood cell lysis. For CD11b⁺ cell isolation, an anti-CD11b antibody (BioLegend, clone M1/70; 5 µL per sample) was added and incubated on ice for 15 min, followed by washing and incubation with SAV beads (BioLegend; 5 µL) for an additional 15 min on ice. After dilution to 500 µL with buffer, CD11b⁺ cells were isolated using MACS MS columns (Miltenyi Biotec). Cells were plated at 1.0 × 10⁶ cells per well in 6-well plates and cultured in 1.5 ml of differentiation medium (DMEM high glucose, 10% FBS, 1× GlutaMAX, 1% penicillin/streptomycin) supplemented with 50 ng ml⁻¹ M-CSF (PeproTech, AF-315-02) and 20 ng ml⁻¹ mouse IL-4 (R&D Systems, 404-ML). Recombinant mouse BST2 (R&D Systems, 9940-BS) was added at concentrations ranging from 0.1 to 1.0 µg ml⁻¹ for stimulation.

#### Flow cytometry analysis of cell proliferation and differentiation in BMDMs

After BST2 stimulation, changes in cell number were assessed using a cell counter, and an effect was observed on day 3. Therefore, cell proliferation was analyzed on day 3. Following the BMDM differentiation described above, the cells were resuspended in 1 × 10⁶ cells ml⁻¹, stained with 1 µM ViaFluor 488 SE Cell Proliferation Kit (Biotium, 99840), and incubated for 10 min at room temperature. The dilution of the fluorescence intensity due to cell division was analyzed by flow cytometry. On the basis of the antibodies listed in the antibody table, the cells were separated into CD11b⁻F4/80⁻ and CD11b⁺F4/80⁺ populations, and ViaFluor 488 fluorescence was evaluated in each fraction. To assess the differentiation markers of BMDMs, the expression levels of CD11c, MHC II (I-A/I-E), CD200, and CX3CR1 were evaluated in 7-AAD⁻ live cells gated as CD11b⁺F4/80⁺Ly6G⁻Ly6C low–negative cells.

#### Bulk RNA-seq for cocultured endothelial cells and BMDMs

Bulk RNA sequencing was performed on adipose endothelial cells subjected to either coculture with macrophages or stimulation with TNFα, as well as on BMDMs treated with recombinant BST2. For the BMDM experiments, treatment with 0.1 µg ml⁻¹ recombinant mouse BST2 was defined as the BST2-low condition, and 1.0 µg ml⁻¹ was defined as the BST2-high condition. The cells were harvested on day 6 after stimulation. Total RNA was extracted using TRIzol™ (Invitrogen, Thermo Fisher Scientific). RNA integrity and concentration were assessed with a 4200 TapeStation system (Agilent, Santa Clara, CA, USA). RNA-seq libraries were prepared using the NEBNext Ultra II Directional RNA Library Prep Kit for Illumina (New England Biolabs, Ipswich, MA, USA) and SPRIselect (Cat. No. B23318, Beckman Coulter, Indianapolis, IN, USA), following the manufacturers’ protocols. Deep sequencing was performed on an Illumina NovaSeq 6000 platform (paired-end mode). The sequenced reads were aligned using Bowtie2 and TopHat (version 1.3.2), and transcriptome assembly was performed with Cufflinks (version 2.0.2).

#### RNA-seq data processing and differential expression analysis

Raw read counts obtained from the HTSeq-count output files were analyzed using the TCC package (version 1.32.0, Bioconductor 3.15) in R (version 4.2.1). Read count matrices were generated for each experimental group and merged according to common gene identifiers. Normalization was performed using the TMM method, and differentially expressed genes (DEGs) were identified with the edgeR test method implemented in TCC. Genes with a false discovery rate (FDR) < 0.05 were considered significantly differentially expressed. For visualization, volcano plots were generated based on the log₂ fold change (m.value) and adjusted p-value (q.value). Genes were classified as having upregulated expression when q < 0.05 and log₂ fold change > 1, downregulated expression when q < 0.05 and log₂ fold change < –1, and nonsignificant otherwise.

#### TPM transformation and heatmap visualization

Gene length data were integrated with the count matrix to calculate transcripts per million (TPM) values. TPMs were calculated by normalizing read counts to gene length and library size using a custom R function. Log₂-transformed TPM values were subsequently scaled by Z score normalization across samples. Heatmaps were generated using the pheatmap package to visualize the expression patterns of genes identified as differentially expressed in the TCC analysis.

#### Gene Ontology (GO) enrichment analysis

Gene Ontology (GO) enrichment analysis was performed for both single-cell and bulk RNA-seq datasets using the clusterProfiler package (version 4.4.4) in R. Differentially expressed genes were converted from gene symbols to Entrez Gene IDs using the org.Mm.eg.db annotation database. Functional enrichment was analyzed with the enrichGO function for the Biological Process (BP) ontology. The Benjamini–Hochberg method was applied to adjust for multiple testing, and terms with an adjusted p-value and q-value < 0.05 were considered significantly enriched. Enrichment results were visualized using enrichplot and ggplot2.

#### C3 depletion

Cobra venom factor-mediated C3 depletion was performed using recombinant cobra venom factor (CVF; A601, Quidel Corporation), provided at 100 µg mL^-1^. CVF was administered intraperitoneally to 15-month-old mice at a dose of 12.5 µg in 125 µL per mouse (n = 4; 3 males and 1 female). Littermate control mice received 125 µL of PBS intraperitoneally. The mice were sacrificed on day 5 after treatment, and the hearts were harvested and prepared for cryosectioning. C3 depletion was confirmed by reduced C3 immunofluorescence signal in heart sections.

#### Recombinant BST2 administration and Oil Red O staining

Atherosclerotic lesion analysis was performed in ApoE knockout mice subjected to cholesterol loading. ApoE knockout mice were fed a high-cholesterol diet (Research Diets, D12079B) starting at 12–15 weeks of age. Recombinant BST2 (Sino Biological, 51043-M07H) was reconstituted in sterile PBS and administered intraperitoneally at a dosage of 5 μg per injection, three times per week. Control mice received sterile PBS as vehicle control using the same injection schedule. The mice were analyzed after 9 weeks of treatment. The study was performed in three independent treatment cohorts, and data were pooled for analysis. After euthanasia, the aorta from the cardiac base to the femoral bifurcation, together with tissue surrounding the aortic valve, was collected. For aortic root analysis, samples from the cardiac base were fixed in 4% paraformaldehyde for 3 h, cryoprotected in 30% sucrose for 24 h, embedded in OCT compound, and sectioned at an 8 μm thickness. Serial cryosections were collected at 100 μm intervals. The section in which the aortic valve leaflets and the sinus of Valsalva were no longer visible was defined as the reference point, and lipid deposition was quantified over a 500 μm region toward the aortic valve. For Oil Red O staining, the sections were air-dried for 30 min, quenched with 100 mM glycine/PBS (pH 7.2) for 10 min, incubated in propylene glycol for 3 min, and stained with Oil Red O solution (Polysciences, 25962-250) for 15 min, followed by differentiation in 85% propylene glycol for 5 min. The nuclei were counterstained with Gill’s hematoxylin for 30 s, mounted with Aqua-Poly/Mount, and imaged under a light microscope. For en face aortic staining, the aorta, including the three aortic arch branches, was carefully dissected and opened longitudinally to expose the luminal surface, followed by Oil Red O staining. The stained aortas were photographed under a stereomicroscope (Nikon) equipped with a digital camera. For en face quantification, the descending aorta from the aortic arch to 1.5 cm distal to the arch was analyzed. The plaque area on the luminal surface was quantified as the percentage of Oil Red O–positive area relative to the total luminal surface area within this region. Quantification of lipid deposition was performed using ImageJ software by calculating the percentage of Oil Red O–positive area relative to the total luminal or lesion-assessed area, as described previously in standard atherosclerosis protocols^67^.

### QUANTIFICATION AND STATISTICAL ANALYSIS

Statistical analyses were performed as indicated in the figure legends. For the ridge plot analyses of the Aging_1 scores across vascular subtypes, comparisons between the two groups were performed using Welch’s t test. For spatial transcriptomic analyses, spatial units were summarized at the sample or predefined tissue-region level for statistical interpretation, whereas bin- or cell-level distributions were used primarily for visualization. Comparisons of the Aging_1 score comparisons between two groups were performed using a two-sided Wilcoxon rank-sum test. For the physical data such as body weight and blood glucose level, and flow-cytometric analyses comparing the two groups, including monocyte numbers and TNF expression, two-sided Student’s t tests were used. En face Oil Red O analysis of the whole aorta and Aortic root Oil Red O analysis were analyzed using a two-sided Student’s t test. For experiments involving multiple groups, one-way ANOVA was used. For the recombinant BST2 stimulation experiments, comparisons between each BST2-treated group and the 0.0 µg ml^-1^ control group were performed using Dunnett’s multiple comparisons test. For image-based quantification experiments involving multiple groups, including C3 depletion experiments, group comparisons were performed using Tukey’s multiple comparisons test after one-way ANOVA. A p value < 0.05 indicated statistical significance.

