## Supplementary material for "Organ-resolved endothelial regulatory programs link aging and metabolic overload to vascular immune remodeling": Document S1. Supplemental figures and legends

**Document S1. Figures S1–S7 and Supplemental Figure Legends**

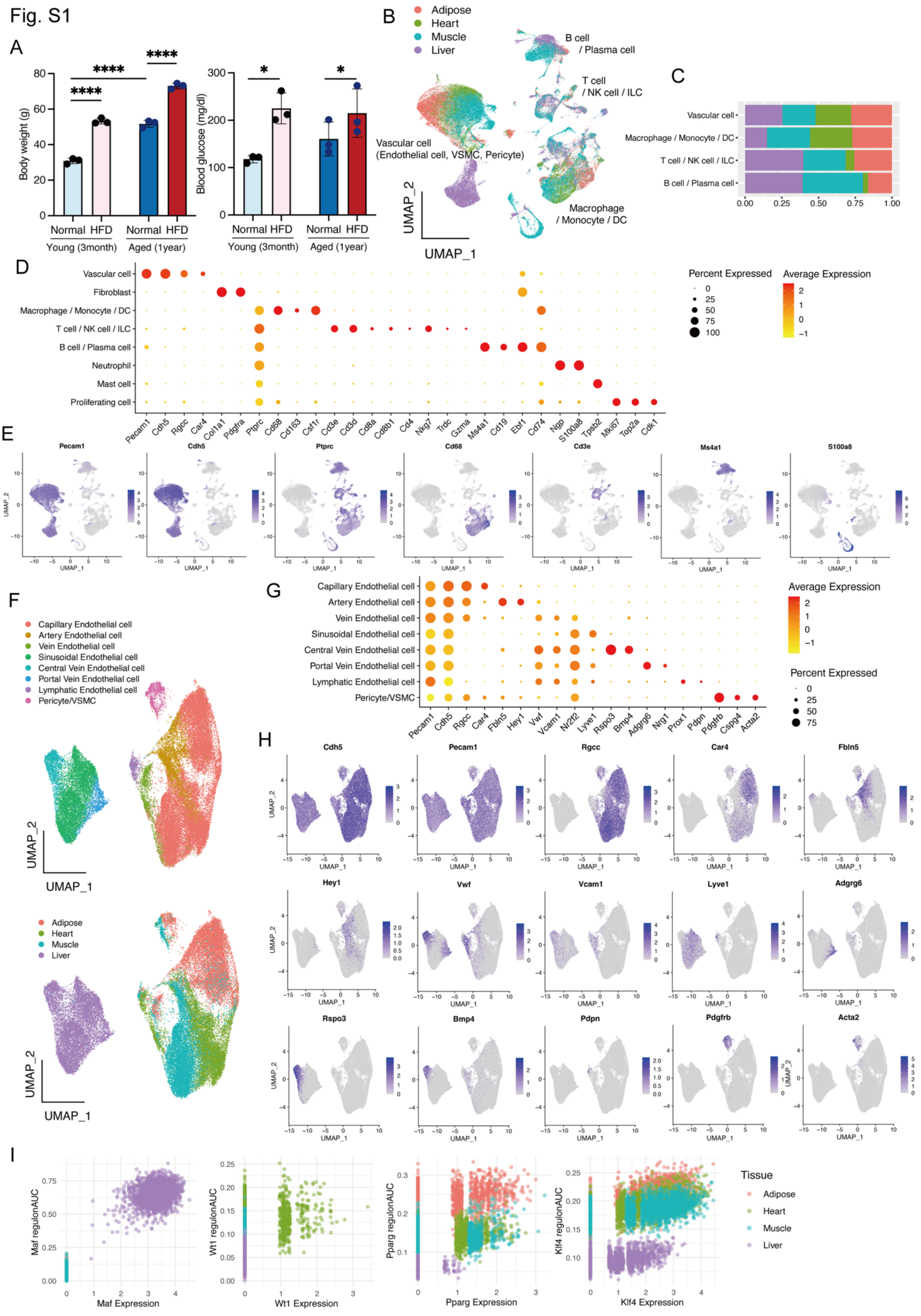


**Figure S1. Sample characteristics and cell annotation for single-cell endothelial analysis, related to Figure 1**

(A) Body weight and ad libitum blood glucose levels at the time of sample collection in each experimental group.

(B) UMAP visualization of cells shown in Figure 1B, colored by organ of origin.

(C) Proportions of annotated cell populations across organs and experimental conditions.

(D) Bubble plot showing representative marker gene expression used for major cell population annotation.

(E) Feature plots showing expression of selected marker genes from (D).

(F) Reclustering of vascular cells used for detailed endothelial and mural cell annotation.

(G) Bubble plot showing representative marker gene expression used for vascular cell subset annotation.

(H) Feature plots showing expression of selected vascular marker genes from (G).

(I) Scatter plot comparing regulon AUC scores with corresponding transcription factor RNA expression in the SCENIC analysis.

Fig.S2

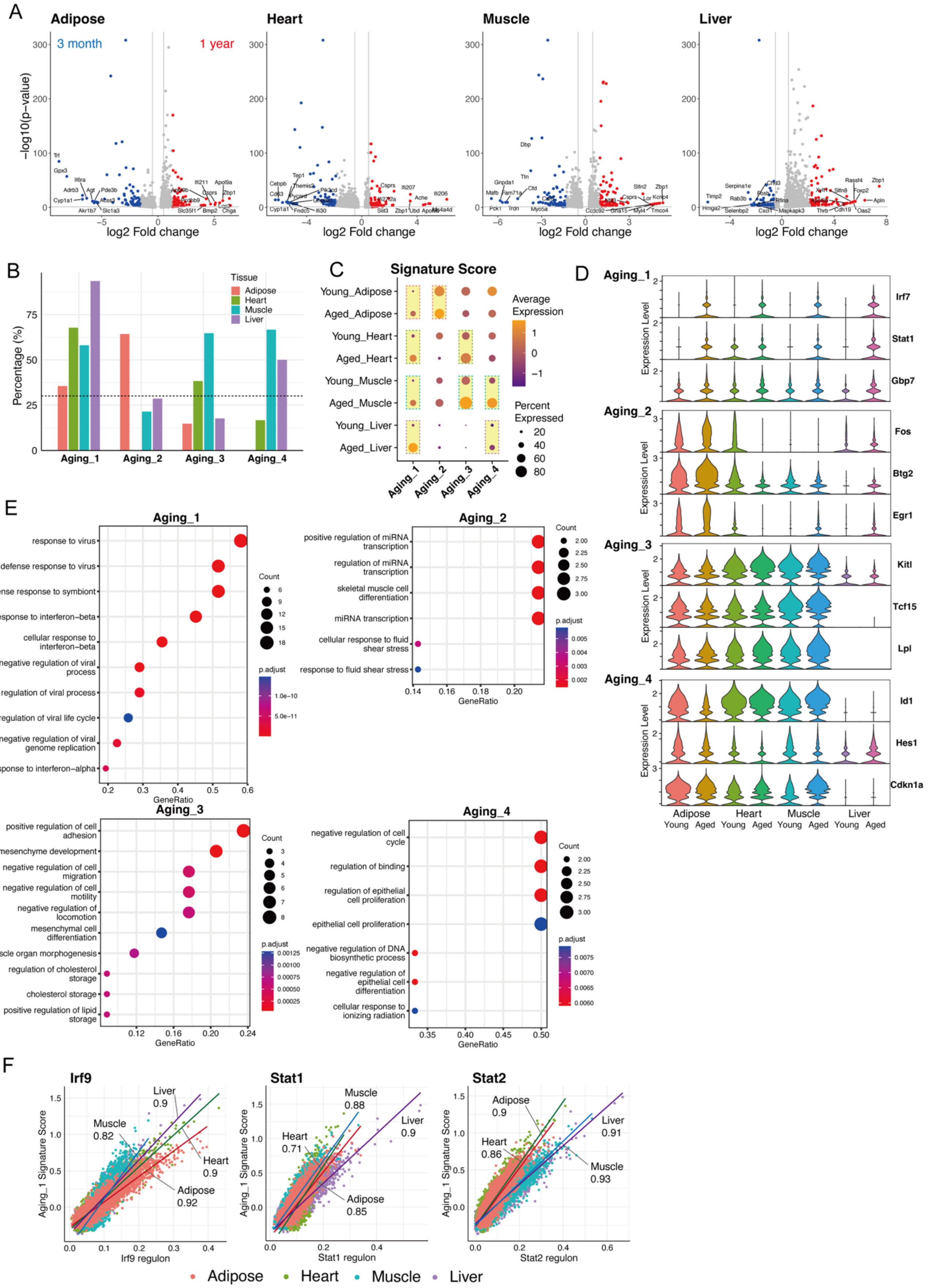


**Figure S2. Supporting analyses of aging-associated endothelial gene programs, related to Figure 2**

(A) Volcano plots showing differentially expressed genes between young and aged endothelial cells in each organ. Genes upregulated in aged endothelial cells were used to select the top 100 aging-associated genes for each organ.

(B) Fraction of genes within each Aging module that were included in the organ-specific top 100 aging-upregulated gene lists.

(C) Signature scores for Aging_1–Aging_4 modules across organs and age groups. Scores were calculated using expression-matched control genes.

(D) Violin plots showing expression of representative genes from each Aging module across organs and age groups.

(E) Gene ontology analysis of Aging_1–Aging_4 gene modules.

(F) Additional single-cell correlation plots between regulon activity and corresponding Aging module signature scores. Each dot represents one endothelial cell; Pearson correlation coefficients are indicated.

Fig.S3

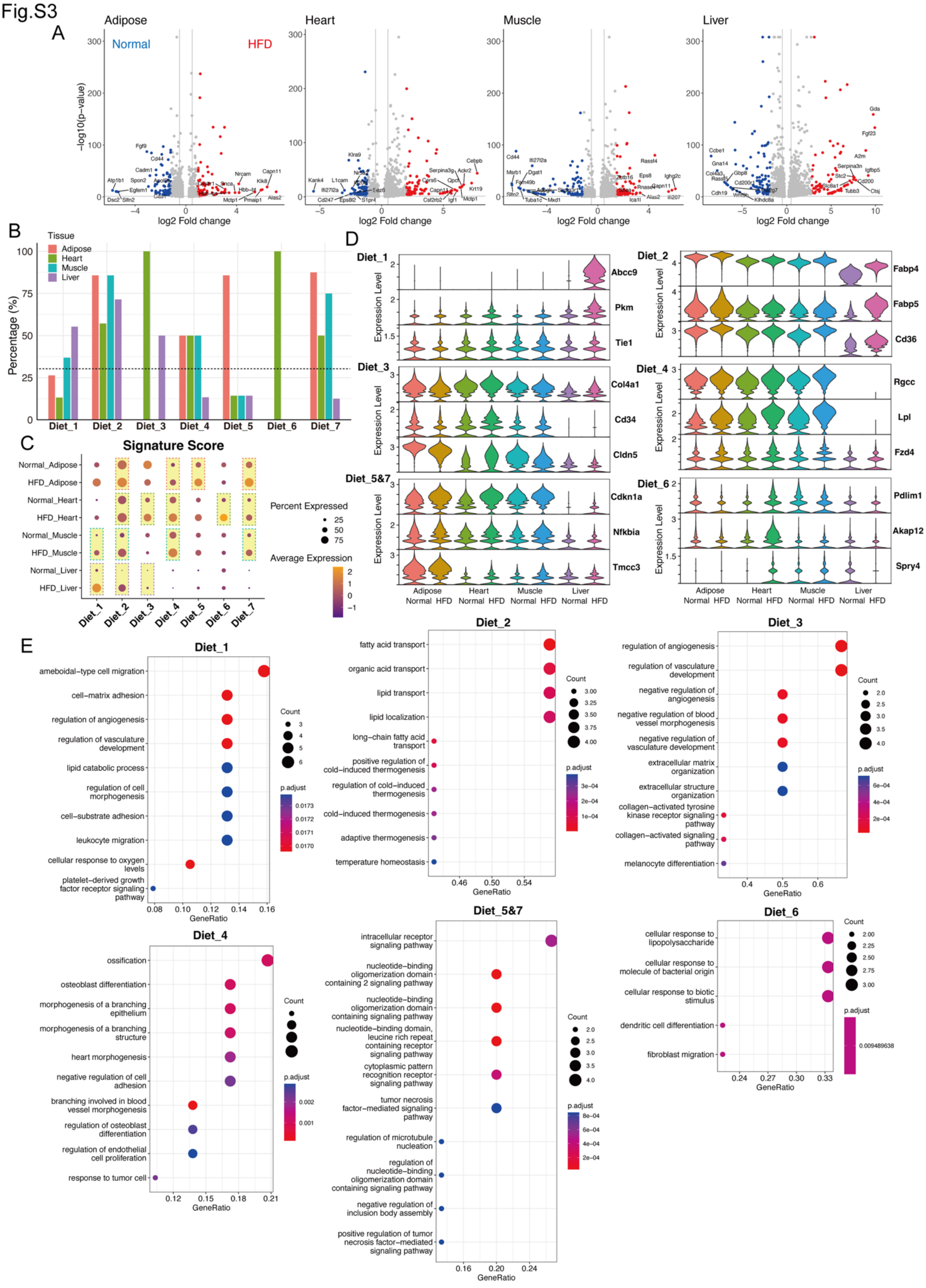


**Figure S3. Supporting analyses of diet-associated endothelial gene programs, related to Figure 3**

(A) Volcano plots showing differentially expressed genes between normal diet and HFD-treated endothelial cells in each organ. Genes upregulated under HFD conditions were used to select the top 100 diet-associated genes for each organ.

(B) Fraction of genes within each Diet module that were included in the organ-specific top 100 HFD-upregulated gene lists.

(C) Signature scores for Diet_1–Diet_7 modules across organs and diet conditions. Scores were calculated using expression-matched control genes.

(D) Violin plots showing expression of representative genes from each Diet module across organs and diet conditions.

(E) Gene ontology analysis of Diet_1–Diet_7 gene modules.

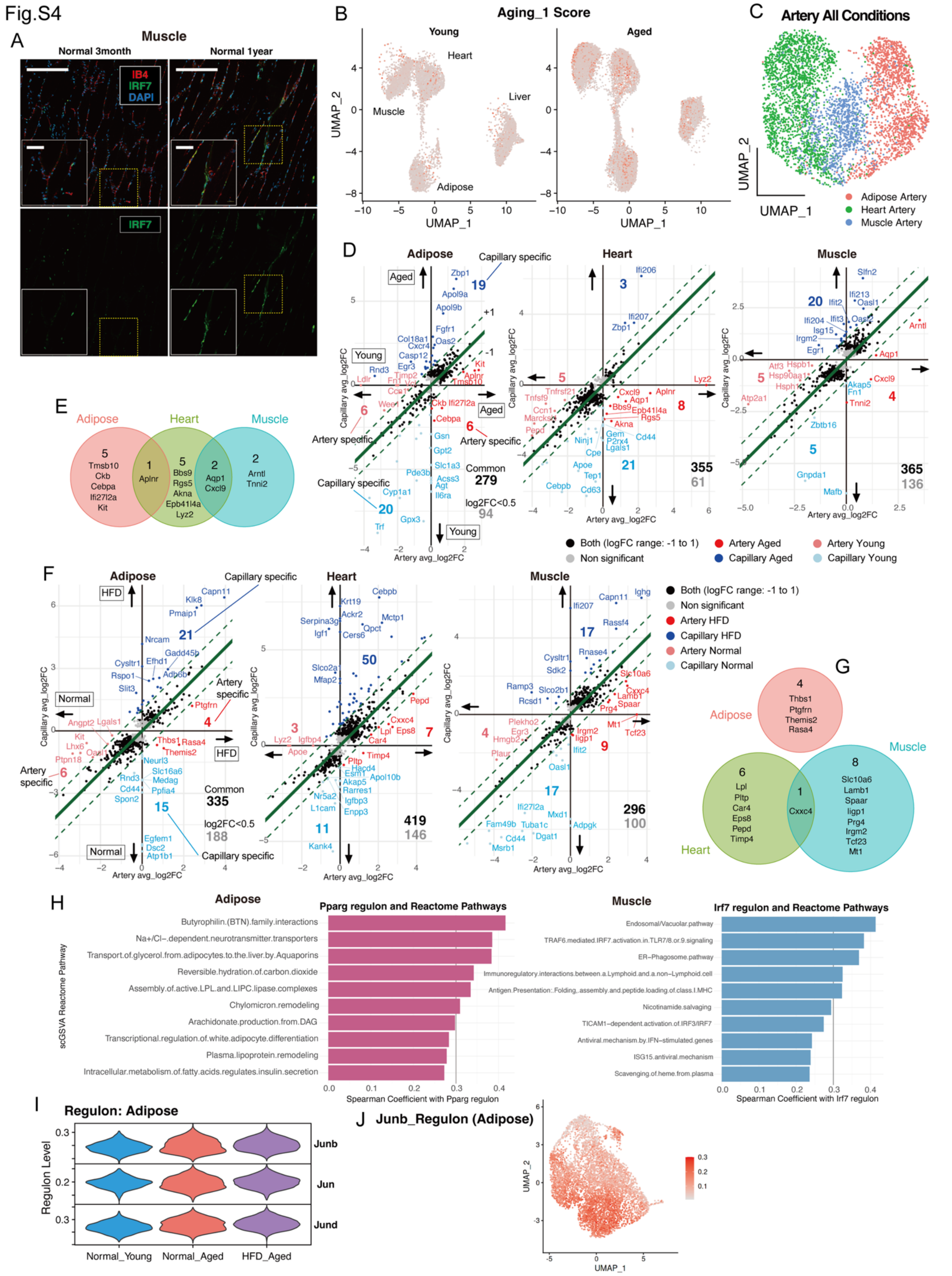


**Figure S4. Supporting analyses of vascular-bed-shared and organ-biased endothelial stress programs, related to Figure 4**

(A) Immunofluorescence images showing IRF7 expression in skeletal muscle from young and aged mice. Endothelial cells were labeled with IB4. Scale bars: low magnification, 200 μm; high magnification, 50 μm.

(B) UMAP projection of Aging_1 scores onto the endothelial cell map shown in Figure 2A. Each point represents one endothelial cell from young or aged mice, and color intensity indicates the Aging_1 score.

(C) UMAP visualization of arterial endothelial cells from adipose tissue, heart, and skeletal muscle across all experimental conditions.

(D) Comparison of aging-responsive genes between arterial and capillary endothelial cells. The green diagonal line indicates the 1:1 relationship, and dashed lines indicate ±1 SD from this line, highlighting genes with relatively artery- or capillary-biased aging responses.

(E) Overlap analysis of the top aging-upregulated genes in arterial endothelial cells from adipose tissue, heart, and skeletal muscle.

(F) Comparison of HFD-responsive genes between arterial and capillary endothelial cells. The green diagonal line indicates the 1:1 relationship, and dashed lines indicate ±1 SD from this line, highlighting genes with relatively artery- or capillary-biased HFD responses.

(G) Overlap analysis of the top HFD-upregulated genes in arterial endothelial cells from adipose tissue, heart, and skeletal muscle.

(H) Representative regulon–pathway correlations identified by scGSVA analysis.

(I) Regulon activity of AP-1 family members in adipose endothelial cells across age and diet conditions.

(J) Reclustering of adipose capillary ECs under normal diet conditions, colored by Junb regulon activity.

Fig.S5

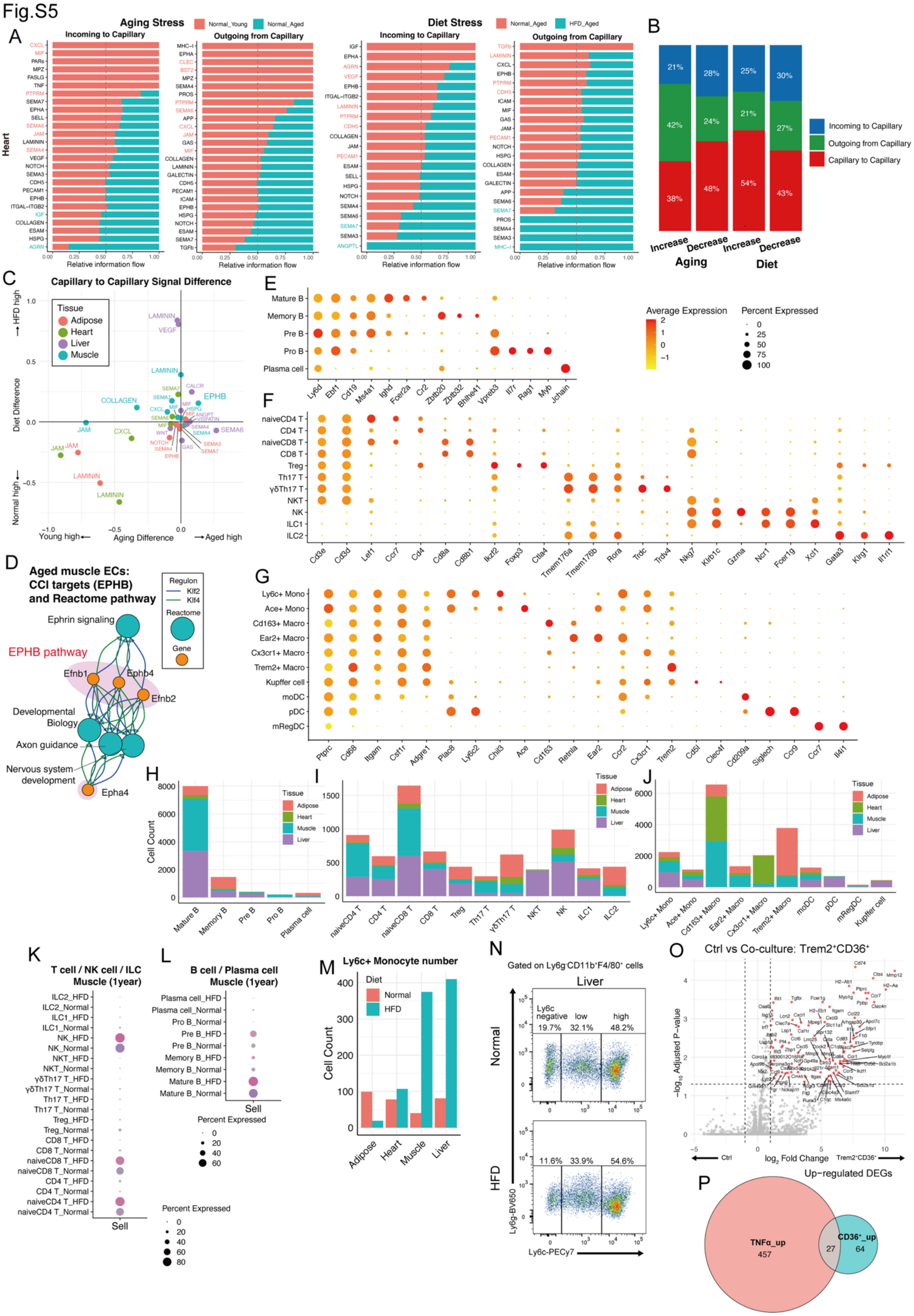

**Figure S5. Supporting analyses of stress-responsive endothelial–immune cell–cell interactions, related to Figure 5**

(A) CellChat rankNet comparison plots showing relative information flow of signaling pathways incoming to or outgoing from capillary endothelial cells under aging or HFD stress. Representative heart datasets are shown.

(B) Fraction of altered signaling pathways classified as capillary-to-capillary, outgoing from capillary endothelial cells, or incoming to capillary endothelial cells. Increased and decreased pathways under aging and HFD stress are shown separately.

(C) Scatter plot comparing capillary-to-capillary signaling differences under aging and HFD stress across organs. Each dot represents a signaling pathway, colored by organ.

(D) Integrated view of EPHB-related cell–cell interaction targets and regulon-pathway associations in aged skeletal muscle endothelial cells. EPHB-associated CCI genes, including *Efnb1*, *Efnb2*, *Epha4*, and *Ephb4*, were compared with Klf2/Klf4 regulon targets and scGSVA-derived Reactome pathways.

(E–G) Dot plots showing marker gene expression used for subclustering immune lineages. B cell lineage subsets are shown in (E), T cell lineage subsets in (F), and Macrophage lineage subsets in (G).

(H–J) Cell counts of immune cell subsets across organs. B cell lineage subsets are shown in (H), T cell lineage subsets in (I), and Macrophage lineage subsets in (J).

(K and L) Sell expression in skeletal muscle immune subsets under normal diet and HFD conditions. T cell lineage subsets are shown in (K), and B cell lineage subsets are shown in (L).

(M) Single-cell RNA-seq-based quantification of $Ly6c^{+}$ monocyte numbers across organs under normal diet and HFD conditions.

(N) Representative flow-cytometry plots showing Ly6c-high monocytes in liver under normal diet and HFD conditions. Cells were gated from $Ly6G^{-}CD11b^{+}F4/80^{+}$ populations.

(O) Volcano plot of bulk RNA-seq differential expression analysis comparing control adipose endothelial cells with endothelial cells co-cultured with $Trem2^{+}CD36^{+}$ adipose macrophages.

(P) Overlap of genes upregulated by $Trem2^{+}CD36^{+}$ macrophage co-culture and genes upregulated by TNFα stimulation in adipose endothelial cells.

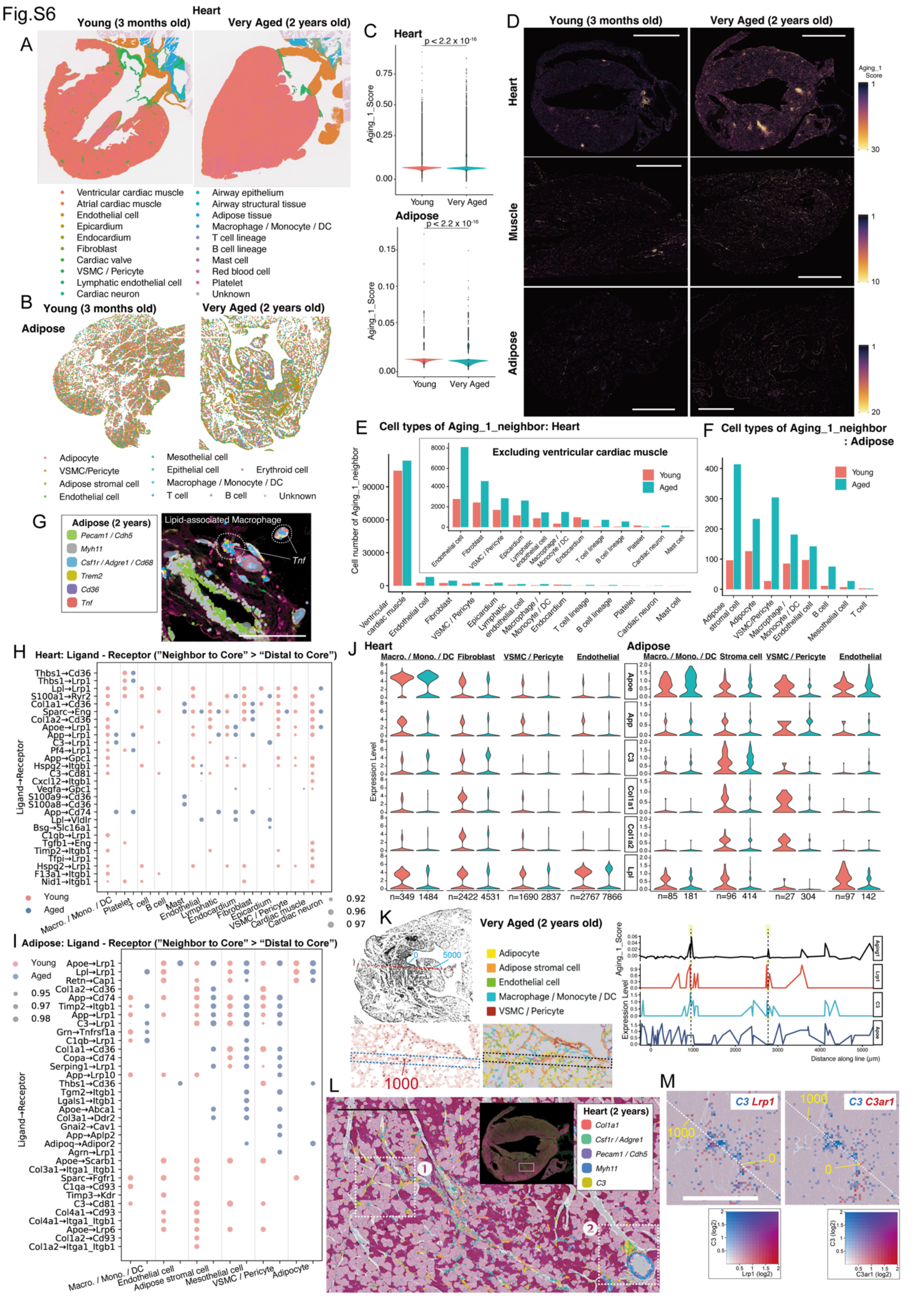


**Figure S6. Supporting spatial transcriptomic analyses of perivascular ISG clusters, related to Figure 6**

(A) Spatial annotation of heart sections from young and very aged mice using Visium HD 8 μm bin data. Cell identities were assigned by label transfer using single-cell reference datasets.

(B) Spatial annotation of adipose tissue sections from young and very aged mice using Visium HD cell segmentation data.

(C) Violin plots showing Aging_1 module scores in spatial bins from heart and segmented cells from adipose tissue in young and very aged mice. Each point represents one bin or segmented cell. P values were calculated at the bin or segmented-cell level and therefore reflect differences across spatial units rather than independent biological replicates.

(D) Xenium in situ gene expression profiling showing Aging_1 scores in heart, adipose tissue, and quadriceps femoris muscle from young and very aged mice. Scale bar, 2 mm.

(E) Cell-type composition of Aging_1 neighbor cells in heart. The inset shows the same analysis after excluding ventricular cardiac muscle.

(F) Cell-type composition of Aging_1 neighbor cells in adipose tissue.

(G) Xenium visualization of a perivascular region in very aged adipose tissue. LAMs were identified as Macrophage lineage cells expressing *Trem2* and/or *Cd36*, and representative *Tnf*-expressing LAMs are indicated by dashed circles. Cell boundaries were displayed according to the Xenium segmentation method: boundary stain method, pink; interior RNA method, yellow; and nuclear expansion method, blue. Scale bar, 100 μm.

(H) LIANA-based ligand–receptor analysis of heart Aging_1 neighbor-to-core signaling. Ligand–receptor pairs with stronger signaling from neighbor to core than from distal to core are shown. Dot size indicates LIANA-derived LRscore.

(I) LIANA-based ligand–receptor analysis of adipose Aging_1 neighbor-to-core signaling. Ligand–receptor pairs with stronger signaling from neighbor to core than from distal to core are shown. Dot size indicates LIANA-derived LRscore.

(J) Expression of the six shared neighbor-to-core ligands identified in Figure 6G within Aging_1 neighbor cells. Violin plots show expression in selected cell populations from heart and adipose tissue. Numbers indicate the number of bins or segmented cells included in each group.

(K) Line-profile analysis across an Aging_1 core–neighbor region in very aged adipose tissue. The line and distance labels indicate position in micrometers from the defined origin.

(L) Xenium visualization of *C3*-expressing regions in the very aged heart. H&E images are overlaid with white cell boundaries defined by Xenium cell segmentation. The region was selected from a nearby section corresponding to the very aged heart area shown in (D) and the ISG cluster analyzed in Figure 6I. Regions ❶ and ❷ indicate the areas magnified in Figure 6L. Scale bar, 200 μm.

(M) Loupe Browser visualization of *C3* with *Lrp1* or *C3ar1* expression in very aged tissue sections. Scale bar, 1 mm.

Fig.S7

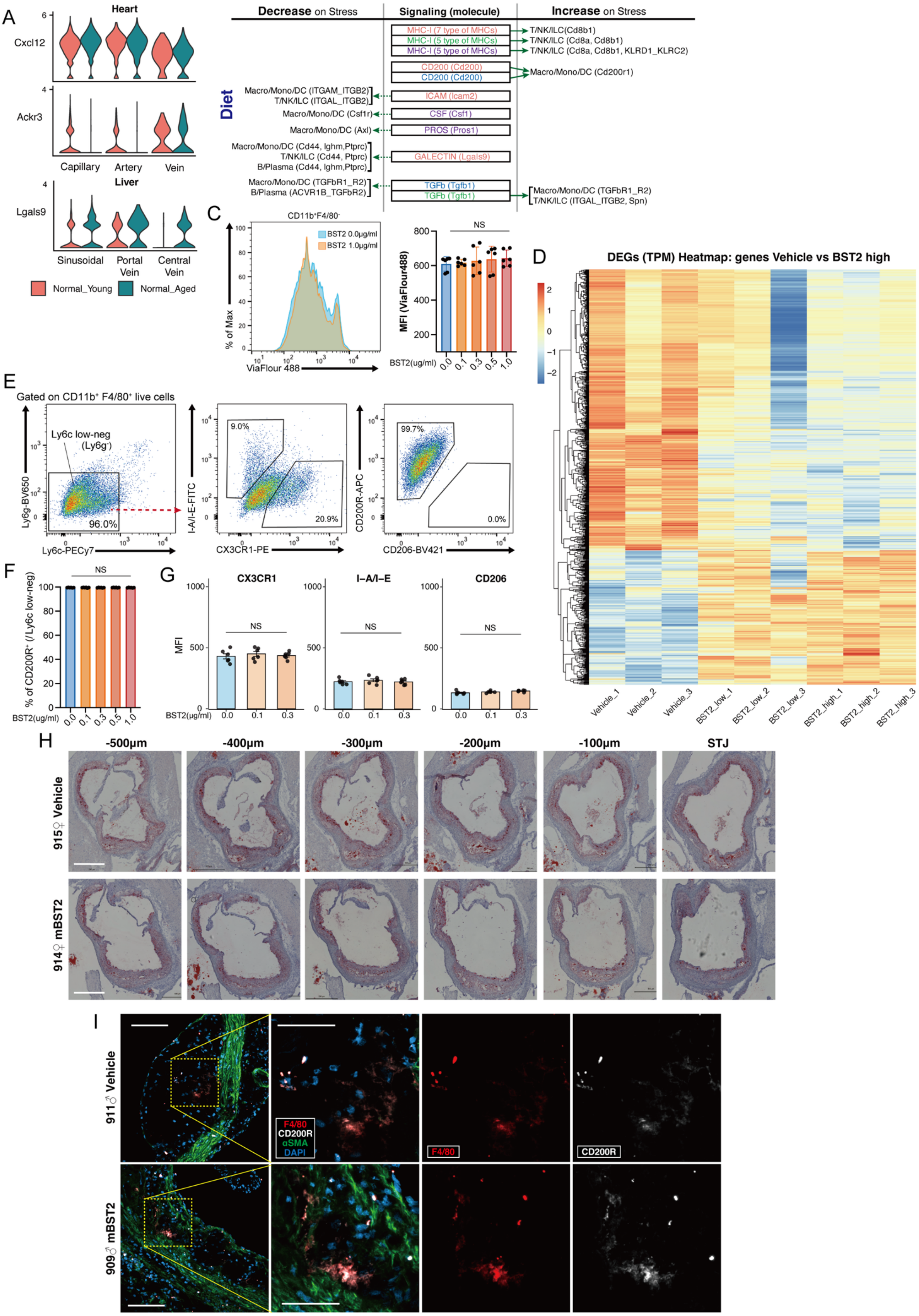


**Figure S7. Supporting analyses of endothelial effector molecules and BST2-mediated macrophage differentiation, related to Figure 7**

(A) Violin plots showing expression of representative CCI-associated endothelial molecules identified in the regulon–effector analysis. *Lgals9* expression is shown in liver endothelial subsets, and *Cxcl12* and *Ackr3* expression are shown in heart capillary, arterial, and venous endothelial cells under young and aged conditions.

(B) Outgoing signaling from capillary and sinusoidal endothelial cells to immune lineages under diet stress. Stress-regulated signaling pathways are shown by organ, sender direction, and recipient lineage.

(C) Cell-proliferation dye dilution analysis using ViaFluor 488 in $CD11b^{+}F4/80^{-}$ cells at day 3 of macrophage differentiation after recombinant BST2 stimulation.

(D) Heatmap showing differentially expressed genes in vehicle, BST2-low, and BST2-high treated cells at day 6 of macrophage differentiation. Treatment with 0.1 μg $ml^{-1}$ recombinant mouse BST2 was defined as BST2-low, and 1.0 μg $ml^{-1}$ as BST2-high.

(E) Flow-cytometry gating strategy for $Ly6G^{-}Ly6C^{low\text{-}negative}$ macrophage-lineage cells used for marker analysis at day 6 of macrophage differentiation.

(F) Quantification of the percentage of $CD200R^{+}$ cells among $Ly6G^{-}Ly6C^{low\text{-}negative}$ macrophage-lineage cells after recombinant BST2 stimulation. NS, not significant.

(G) Flow-cytometric quantification of CX3CR1, I-A/I-E, and CD206 expression in $Ly6G^{-}Ly6C^{low\text{-}negative}$ macrophage-lineage cells after recombinant BST2 stimulation. NS, not significant.

(H) Representative aortic root sections from vehicle- and recombinant BST2-treated ApoE-deficient mice. Numbers indicate distance in micrometers from the sinotubular junction toward the base side of the aortic root. STJ: sinotubular junction. Scale bars, 500 μm

(I) Representative immunofluorescence staining of aortic root sections from the mice analyzed in Figure 7N. Sections were stained for F4/80, CD200R, αSMA, and DAPI. Magnified views correspond to the boxed regions in the overview images. Scale bars, 100 μm in overview images and 50 μm in magnified views.
